# Delineation of a thrombin receptor-stimulated vascular smooth muscle cell transition generating cells in the plaque-stabilising fibrous cap

**DOI:** 10.1101/2024.07.02.600985

**Authors:** James CK Taylor, Matthew D Worssam, Sebnem Oc, Jordi Lambert, Krishnaa T Mahbubani, Kirsty Foote, Allie Finigan, Yee-Hung Chan, Nichola Figg, Murray C H Clarke, Martin R Bennett, Helle F Jørgensen

## Abstract

**Aims:** Vascular smooth muscle cells (VSMCs) accumulate in atherosclerotic plaques and exhibit remarkable phenotypic plasticity, contributing to both plaque growth and stability. The plaque-stabilising fibrous cap is rich in VSMC-derived cells, yet the cellular transitions and regulatory mechanisms governing fibrous cap formation remain unclear. We aimed to delineate the VSMC phenotypic transitions associated with this critical process.

**Methods and Results:** Mapping of lineage-traced VSMCs during plaque development revealed investment of VSMCs prior to fibrous cap formation. Using single-cell RNA-sequencing (scRNA-seq) profiles of lineage-traced VSMCs from atherosclerotic and acutely injured mouse arteries, we identified a disease-specific VSMC state co-expressing contractile genes with extracellular matrix (ECM) components (including fibrillar collagens and elastin) and NOTCH3, which are associated with fibrous cap formation. Computational trajectory analysis predicted that this proposed fibrous cap-related VSMC (fcVSMC) state arises from a previously described plastic, intermediate VSMC population expressing SCA1 and VCAM1. Clonal analysis further showed that NOTCH3^+^ fcVSMCs derive from intermediate VSMCs in both atherosclerosis and an acute vascular injury model, suggesting a conserved disease-relevant mechanism. The fcVSMCs were enriched in plaque fibrous caps compared to lesion cores, consistent with a role in fibrous cap formation. By combining scRNA-seq trajectory analysis and spatial transcriptomics of human atherosclerotic plaques, we identified protease-activated receptor-1 (PAR1) as a candidate regulator of fcVSMC generation. PAR1 was expressed by VSMCs in human plaque fibrous caps and, PAR1 activation by thrombin induced expression of contractile genes and ECM components associated with the fcVSMC state in human VSMCs.

**Conclusions:** Our findings identify a VSMC transition linked to fibrous cap formation in atherosclerosis and show this is modelled by vascular injury. We identify VSMC-expressed PAR1 as a potential therapeutic target for promoting plaque stability by driving the transition to the matrix-producing, fibrous cap-associated VSMC state.

## 1. Introduction

Vascular smooth muscle cell (VSMC) accumulation is a hallmark of atherosclerosis^1-3^. VSMCs are also the main cell-type responsible for neointima formation following vessel occlusion and injury^4-6^. In health, VSMCs are typically found in the medial arterial layer, where they are quiescent and express components of the contractile machinery (such as *MYH11, CNN1 and ACTA2*) to control blood pressure and flow^7,8^. However, these cells retain remarkable plasticity and can dedifferentiate in atherosclerosis and in response to injury or inflammation. This involves downregulating the contractile apparatus and becoming migratory, proliferative, substantial producers of extracellular matrix (ECM), and the cells can acquire a range of different phenotypes^1,2,9^. In atherosclerotic plaques, VSMCs generate cells that make the plaque-stabilising fibrous cap, as well as cells in the lesion core that have characteristics of mesenchymal cells, phagocytes, and osteochondrocytes, which may contribute to lesion destabilisation^2,4,10,11^. Delineating the governance of these distinct VSMC-derived cells in disease will therefore aid understanding of the functional role of these cells in disease.

Multi-colour lineage-tracing studies in disease models have shown that VSMC-contribution to atherosclerotic plaques and injury-induced neointimal lesions is oligoclonal, whereby one or very few pre-existing VSMCs give rise to all lesional VSMCs^12-16^. By extension, the clonal progeny of individual VSMCs can generate the diverse phenotypes observed in atherosclerotic lesions^12,17^. In mice, VSMCs with reduced levels of contractile markers that express stem cell antigen 1 (SCA1, encoded by *Ly6a*) have been suggested to represent a plastic VSMC state capable of giving rise to diverse plaque VSMC phenotypes^17^. Humans, which do not have a SCA1/*Ly6a* homologue, also have VSMC populations with an equivalent transcriptional profile to mouse SCA1+ VSMCs^18^. This plastic VSMC state is rarely detected in healthy arteries and is induced upon VSMC dedifferentiation in response to disease stimuli^17,18^. In atherosclerosis, such cells showed substantial overlap in the transcriptional signature of ‘activated’ VSMC-derived cells expressing osteochondrocyte and phagocyte markers^17^. Computational inference of VSMC transdifferentiation pathways^19^, functional *in vitro* experiments with isolated SCA1-expressing VSMCs^20^ and use of a dual lineage tracing system^21^ have all provided substantial evidence that VSMCs co-expressing SCA1, VCAM1 and LGALS3 act as intermediates and give rise to the diverse VSMC states observed in atherosclerotic plaques. Accordingly, this “intermediate modulated” VSMC (imVSMC) population could represent a novel therapeutic target, whereby its disease-induced transdifferentiation to diverse cell states could be biased towards those states which are beneficial to disease outcome.

The fibrous cap is rich in VSMCs, many of which express typical contractile VSMC markers, as well as ECM components such as fibrillar collagens and elastin^2,22,23^. This structure is imperative to mitigating plaque rupture and downstream cardiovascular events. Previous work has highlighted several candidate factors involved in promoting the formation of a VSMC-rich fibrous cap, such as PDGFRB^24^, TCF21^25^ and retinoic acid^20^. Compelling evidence has also been presented that suggests, although loss of NOTCH3 signalling characterises VSMC lesion investment, subsequent reactivation is critical in the formation of a fibrous cap with contractile marker-expressing VSMCs^26^. However, the VSMC states underpinning fibrous cap formation and how cap formation is related to the imVSMC state remains to be elucidated.

Here, we identify a transcriptional VSMC state in atherosclerosis that we propose underlies fibrous cap formation. Further, we find that VSMCs in this state are generated from cells in the imVSMC state. Interestingly, this VSMC state transition is present in both atherosclerotic and acutely injured arteries, suggesting importance for general VSMC regulation. We then identify candidate regulators of this fibrous cap-associated transition and highlight VSMC-expressed thrombin receptor as a candidate driver of fibrous cap formation.

## 2. Methods

### Animals and procedures

All experiments were carried out according to the UK Home Office regulations under PPLs P452C9546 and PP7513347 and were approved by the University of Cambridge Ethical Reviews Committee. VSMC labelling was achieved using Myh11-CreER^T2^; Rosa26-Confetti (Myh11^Confetti^) and Myh11-CreER^T2^; Rosa26-eYFP (Myh11^eYFP^) mice aged 6-8 weeks by 10 intraperitoneal injections of 1 mg tamoxifen (Sigma) dissolved in corn oil over 2 weeks. Animals were rested for at least 7 days before inducing disease. Surgery was performed as described previously^12^. Briefly, Myh11^Confetti^ and Myh11^eYFP^ animals were subcutaneously administered pre-operative analgesic (∼0.1 mg/kg body weight, buprenorphine), anaesthetised with 2.5-3% isoflurane by inhalation (1.5 L/min) and the left carotid artery tied off with 6-0 silk suture. Lineage-labelled Myh11^Confetti^; *Apoe^-/-^* mice were fed a high fat diet (HFD, Special Diets Services, containing 21% fat and 0.2% cholesterol) for the length of time indicated. Animals were culled at the indicated timepoint after surgery by CO_2_ asphyxiation and perfused with phosphate-buffered saline (PBS) prior to tissue removal. Mouse arteries were dissected free from adipose and connective tissue before experiment-specific processing (as described below).

### Single-cell RNA-sequencing (scRNA-seq) data generation

Datasets of lineage-labelled VSMCs from healthy and injured mouse carotid arteries were generated as described^18^ after pooling cells from 5 Myh11^eYFP^ animals 11 days post-ligation or 2 control Myh11^eYFP^ animals that did not undergo surgery. After carotid artery harvest and dissection, arteries were incubated for 6 minutes in collagenase type IV (1 mg/mL, Life Technologies) and porcine pancreatic elastase (1 U/mL, Worthington) in DMEM. After removing the adventitia, the medial samples from each animal cohort were pooled and digested in collagenase type IV (2.5 mg/mL, Life Technologies) and porcine pancreatic elastase (2.5 U/mL, Worthington) in DMEM for 90 minutes. Medial cell suspensions were filtered through a 40 µm filter and singlet cells expressing eYFP isolated by fluorescence-activated cell sorting (FACS, Aria-Fusion flow cytometer, BD Bioscience) into PBS with 0.05% BSA and 5 µL/mL recombinant RNAse inhibitor (Sigma). Isolation of single cells was verified with a haemocytometer. Sorted cells (10,000) were pelleted, resuspended in 47 µL of PBS with 0.05% BSA and loaded for Drop-Seq using the 10X Chromium system. Amplified cDNA libraries were sequenced on a HiSeq 4000 system.

### scRNA-seq data analysis

Raw sequencing reads were processed and aligned to the GRCm38 mouse genome (+eYFP cDNA/ORF sequence) through the 10X Genomics Cell Ranger pipeline (v3.1.0). Datasets for Myh11 promoter-driven, lineage-labelled VSMCs in a mouse atherosclerosis model have been published previously^20^ and gene-count matrices were downloaded from the gene expression omnibus (GEO, accession number GSE155513). All datasets were analysed using the R or Python programming language. Key package version numbers, dataset-specific quality control (QC) parameters, normalisation method and numbers of principal components (PCs) used for the analysis are indicated in Supplementary Table 1. For the mouse atherosclerosis dataset, QC parameters defined in the original publication^20^ were used. For cells that passed the QC criteria, expression data was normalised using either the “NormalizeData” or “SCTransform” (described previously^27^) function in Seurat^28^. The major principal components (PC’s), identified using a combination of the Elbow method and “JackStraw” function, were used for Louvain clustering and Uniform Manifold Approximation and Projection (UMAP). The resolution was validated by assessing cluster boundaries using the “sleepwalk” package^29^ (CRAN). Differential gene expression between clusters was determined using the “FindAllMarkers” function in Seurat. Significance was determined by Wilcoxon rank sum test [FDR-adjusted P-value (P-adj)<0.05] and differentially expressed gene thresholds of log_2_ fold-change (log_2_FC)>0.5 and minimum 25% of expressing cells within a cluster. Integration of healthy mouse carotid and atherosclerosis datasets was performed with the standard Seurat workflow using the “FindIntegrationAnchors” and “IntegrateData” functions (first 30 PCs used).

The Slingshot package^30^ (Bioconductor) was used for atherosclerosis and injury dataset trajectory analyses. For this, the Seurat object was transformed to a SingleCellExperiment (SCE) object using the SCE package^31^ (Bioconductor) and trajectory path inference performed with the “slingshot” function using the same number of PCs used for dimensionality reduction and clustering. The cell cluster showing the highest contractile VSMC gene expression was selected as the origin for the analysis. Differential expression across the trajectory path of interest was performed with Generalised Additive Modelling (GAM) using the “gam” package^32^ (CRAN). The 3000 highly variable genes (HVGs) were used as input for this analysis. A GAM (with a Loess term for pseudotime) was fitted for each gene (using the “gam” function) and significance for pseudotemporal-dependent expression assessed (FDR-corrected P-adj<0.05). A magnitude threshold for the difference between the modelled maximum and minimum values in a log scale over trajectory pseudotime of logFC>0.25 was then applied for genes with significant pseudotime-dependent expression. The genes that passed these criteria were grouped into gene “clades” with similar expression patterns based on Pearson correlation distance and the complete linkage method, which were plotted in a heatmap over trajectory path pseudotime (all performed using the “pheatmap” package^33^ (CRAN). The optimal number of clades was determined using the Elbow method.

Partition-based graph abstraction (PAGA^34^) was performed on the injury dataset using functions available within Scanpy^35^ for Python. The package “anndata2ri”^36^ and associated extensions were used to interface R and Python languages. This was used to convert the injury dataset SCE (R) object into an AnnData (Python) object. From here, dimensionality reduction was performed as before using equivalent functions in the Scanpy package. The cluster labels obtained in the original Seurat analysis were retained. PAGA was then run using the “sc.pl.paga” function and an edge connectivity threshold of 0.35 was the highest value that permitted a connected structure of the data. The “sc.pl.paga_compare” function was used to visualise cell and cluster connectivity projections over UMAP plots generated with initial PAGA positions taken as additional input.

### Gene ontology analysis

Gene ontology (GO) analysis of genes differentially expressed between clusters or along trajectory path pseudotimes was carried out using the gProfiler^37^ web tool (https://biit.cs.ut.ee/gprofiler/gost) for all terms against mus musculus or homo sapiens genes (as appropriate) with a term size<1000. The Circos plot was generated using the GOplot package (CRAN).

### Immunofluorescence

Mouse arteries were fixed in 4% formaldehyde (Sigma) for 20 minutes at room temperature (RT), cryopreserved in 30% sucrose in PBS overnight, adjusted for 1 hour in a 50:50 solution of 30% sucrose solution: Optimal Cutting Temperature (OCT) compound (VWR), followed by 1 hour in 100% OCT and snap-frozen in OCT using dry ice. Sections (14 μm for immunostaining, 100 μm for lesional VSMC infiltration analysis) were cut onto Superfrost® Ultra slides (Thermo) and either mounted directly or stained in a humidity chamber protected from light. For immunostaining, cryosections were rinsed in PBS and permeabilised in 0.5% (v/v) Triton X-100 (Sigma Aldrich) for 20 minutes in PBS at room temperature, incubated for 1 hour at RT in blocking buffer [1% (w/v) bovine serum albumin and 10% (v/v) normal goat serum, Dako] before incubation with primary antibodies for NOTCH3 (Abcam, ab23426, 10 μg/mL), VCAM1 (BioLegend, 105702, 10 μg/mL) or isotype control antibodies diluted in blocking buffer overnight at 4°C. Primary antibodies were removed with 3 x 5 minute PBS washes before incubation with Alexa Fluor 647 and Alexa Fluor 750-conjugated secondary antibodies (diluted in blocking buffer) for 1 hour at room temperature. Sections were washed 2 x 5 minutes in PBS, nuclei stained with DAPI (1 μg/mL) for 10 minutes at room temperature, before rinsing in PBS and mounting in RapiClear 1.52 (Sunjin Lab).

### Confocal imaging

Confocal imaging was performed using an SP8 (VSMC infiltration analysis) or Stellaris (immunostaining analysis) laser scanning microscope (Leica) with a 20x lens in sequential, resonant, tile scan mode with 3 μm distance between Z-stack sections. Laser lines and detector settings were selected to avoid spectral overlap; VSMC infiltration analysis (SP8): sequence 1 (405/410-462, DAPI), sequence 2 (458/462-483, CFP and 555/565-632, RFP), sequence 3 (514/520-558, YFP), and sequence 4 (488/490-507, GFP); seven-colour immunostaining (Stellaris): sequence 1 (405/410-440, DAPI and 638/643-710, Alexa Fluor 647), sequence 2 (448/453-475, CFP and 730/735-850, Alexa Fluor 750), sequence 3 (488/493-509, GFP), sequence 4 (514/519-540, YFP) and sequence 5 (561/566-610, RFP). Data was acquired at an optical section resolution of 1024 x 1024 and tiles were stitched using the mosaic merge function in LASX software (Leica). Image analysis was done using Imaris software (9.1.2) to adjust brightness and contrast; isotype control and primary antibody-stained sections were subject to identical manipulations.

### Scoring of lineage-traced VSMC sections

Quantification of VSMC infiltration into plaques was done using 100 μm plaque sections analysed previously in the assessment of VSMC clonality^18^. A total of 155 plaques from the ascending and descending aorta, aortic arch and carotid arteries of 7 animals (4 at week 6.5 and 3 at week 11) were imaged. Plaques with suboptimal clearing (5) were excluded from the analysis, hence a total of 150 were included in the analysis. Quantification of VCAM1 and NOTCH3 staining in plaques was done after immunostaining of 14 μm sections from a subset of plaques (n=16) from animals after 11 weeks HFD (n=3). In total, 26 VSMC clones were identified in these lesions. VSMCs were scored as "luminal edge" when present within three cell-widths of the endothelial boundary or if elongated and part of a luminal clone, "internal elastic lamina (IEL) adjacent (IEL-A)” when inside the plaque but in direct contact with the IEL, and "core" if meeting neither of these criteria. Cap presence was scored when contiguously arranged cells with elongated nuclei (based on DAPI signal) were present at the luminal edge. Elastic lamina breaks were scored in lesions where image quality allowed detection of localised absence of autofluorescence from the internal elastic lamina (IEL). Quantification of VCAM1 and NOTCH3 staining in injury was performed in 3 control vessels or diseased regions of vessels from 3 injured animals per timepoint. Expression of VCAM1 and/or NOTCH3 by individual VSMCs was confirmed by navigating through multiple *Z-*planes.

### Human tissue

Anonymised human artery samples were obtained from patients undergoing carotid endarterectomy, cardiac transplant, valve replacement or by donation after circulatory death. All human tissue acquisition was done under informed consent using protocols approved by the Cambridge or Huntingdon Research Ethical Committee.

### Spatial transcriptomics

A plaque-containing aortic sample from a 54 year old male organ donor was snap-frozen by fully submerging in an isopentane bath cooled by dry ice until frozen, embedded in OCT (2-8°C), sectioned and analysed using VISIUM technology (cat 1000184) according to manufacturer’s instructions (10X). Sequencing data was aligned to a 10X-supplied human reference genome (GRCh38-3.0.0) using Space Ranger (v.1.0). Downstream analysis was performed with R (v.4.0.3). The Spaniel R package^38^ (v.1.2.0) was used to import the data into SingleCellExperiment objects and was also used to create spot images. Any spots with fewer than 200 UMIs were removed. The SingleCellExperiment objects were then converted into SeuratObjects for downstream analysis using Seurat (v.3.1.3). Individual replicates were normalised using the “NormaliseData” function followed by the identification of the top 200 highly variable genes using the “FindVariableFeatures” function. Replicates for each sample were integrated using the FindIntegrationAnchors method, with dims = 1:20 and k.filter = 50. Following this, the data was processed with the following functions: data was scaled with ScaleData, RunPCA was used with npcs = 30, RunUMAP with dims = 1:15 and FindNeighbors with k.param = 10. The FindClusters function was used to cluster spots with a resolution of 0.4. The Seurat function FindAllMarkers was used to identify marker genes for each cluster. All other parameters were selected as default unless otherwise stated.

### Immunohistochemistry

Human arteries were formaldehyde-fixed and paraffin-embedded (FFPE) and sections (4 μm) were dewaxed, processed for antigen retrieval and stained as described^18^. Sections were co-stained for αSMA (DAKO, M0851, 1:400), detected with biotin-coupled anti-Mouse (DAKO, E0433) using Vectastain avidin-coupled alkaline phosphatase with Blue AP substrate solution (Vector Labs), and PAR1 (Abcam, ab32611, 1:200), that was detected with HRP-conjugated anti-Rabbit (Cell Signaling Technology, 8114, using 3,3’-diaminobenzidine tetrahydrochloride (DAB) as peroxidase substrate, SignalStain).

### Cell culture

Human VSMC (hVSMC) cultures were generated from aortas of patients undergoing cardiac transplant or aortic valve replacement. After manually removing the endothelial layer and adventitia, the medial layer was cut into 2-3 mm² pieces, placed into 6-well plates containing 1 ml media [DMEM supplemented with 20% fetal calf serum (FCS),100 U/mL penicillin, 100 µg/mL streptomycin] and cultured to allow cells to migrate out of the tissue (1-2 weeks). After establishment, cells were cultured in hVSMC-specific medium (Promocell, SMC-GM2) supplemented with 100U/mL penicillin, 100µg/mL streptomycin and were studied at passages 2–10. Thrombin treatments were performed in serum-free media. To transiently silence PAR1, hVSMCs were transfected with 50 nM human PAR1-specific siRNA (ON-TARGETplus® SMART Pool, Dharmacon, L-005094-00-0005) or non-targeting control siRNA (ON-TARGETplus® Control Pool, nontargeting pool, Dharmacon, D-001810-10-05) for 72 hours using Lipofectamine RNAiMAX transfection reagent (Invitrogen). Cells were then washed prior to 24 hour thrombin treatments, followed by downstream analysis.

### Bulk expression analysis

Total RNA was isolated using the RNeasy Mini kit (Qiagen), reverse transcribed using QuantiTect Reverse Transcriptase (Qiagen), and cDNA corresponding to 10 ng RNA was analysed using quantitative, real-time, reverse transcription PCR (RT-qPCR) with SsoAdvanced Universal SYBR Green Supermix (Biorad, primer sequences listed in Supplementary Table 2). HMBS was used as a housekeeping gene for data normalisation. Bulk RNA-seq was conducted with RNA isolated from 3 different female hVSMC isolates following a 24 hour thrombin treatment in serum-free medium. Libraries were prepared from oligo-dT-purified mRNA and sequenced on an Illumina Novaseq 6000 (150 bp paired end reads). Raw data reads were trimmed with Trim Galore (v.0.6.7), and aligned to the human genome (GRCh38) with Kallisto (v.0.46.2). Trimmed mean of M-values (TMM) normalization was then conducted. Differential expression analysis was performed using the “voom” normalisation workflow of the limma package^39^ (Bioconductor) with cell line donor as a covariate. Linear models were fitted to the normalised data, and empirical Bayes moderation was applied to the standard errors of the estimated log-fold changes. Contrast matrices were specified to identify differentially expressed genes between treatment groups. An FDR-corrected P-adj<0.05 was considered statistically significant. The thrombin-upregulated gene module for expression analysis in the scRNA-seq data was generated by converting human gene orthologues to mouse using the gProfiler web tool and using these orthologues with the “AddModuleScore” function in Seurat.

To analyse PAR1 protein expression, whole cell protein lysates were prepared in RIPA buffer freshly supplemented with proteinase inhibitors (Millipore) and phosphatase inhibitors (Millipore). Protein concentration was determined using the BCA method (23227, Pierce BCA protein assay kit, Thermo Fisher). Immunoblotting was performed according to standard conditions, using gradient (4-12%) polyacrylamide gels, methanol-based wet transfer and chemiluminescence detection (Amersham ECL detection reagent, GE Healthcare). Primary antibodies for PAR1 (Abcam, ab32611, 1:2000), and GAPDH (loading control, Cell Signaling Technologies, 2118, 1:4000) were detected using HRP-labelled secondary antibodies: goat-anti-rabbit (7074S, Cell Signaling Technology).

### Data availability

The bulk and single cell RNAseq datasets generated in this study have been deposited to the Gene Expression Omnibus (GEO) repository (accession numbers available upon publication). The scRNA-seq datasets from VSMC-lineage-labelled plaque cells from HFD-fed animals are available from GEO, GSE155513. Source data is available upon request.

### Statistical analysis

Statistically significant differences between treatment groups in the RT-qPCR analysis were assessed using the Mann-Whitney test with Bonferroni multiple testing correction. Group sizes represent samples from different donors, different mice, plaques or clones as specified, rather than technical replicates. An adjusted P-value (P-adj) less than 0.05 was considered statistically significant. For assessing the overlap of gene sets, the Chi-squared test was used with a P-value less than 0.05 considered statistically significant. Statistical testing in analysis of single cell and bulk RNA-seq datasets is provided above.

## 3. Results

### 3.1. Plaque VSMC infiltration leading to generation of the fibrous cap

To understand the kinetics of fibrous cap formation, we analysed plaques in arteries of *Apoe^-/-^* mice carrying a Myh11-driven, tamoxifen-inducible Cre recombinase and the Rosa26-Confetti reporter (Myh11^Confetti^), such that stochastic expression of one of four fluorescent proteins is induced by tamoxifen treatment specifically in VSMCs. Following tamoxifen treatment to induce mosaic VSMC labelling, the animals were fed a high fat diet (HFD) for 6.5 (W6.5) or 11 weeks (W11), to trace the clonal progeny of existing VSMC-lineage cells in developing plaques. We found that the proportion of lesions with longitudinally arranged cells adjacent to the endothelium at the luminal edge, indicating formation of a fibrous cap, increased with both time of HFD and plaque size (Figure 1A-C). This is consistent with a correlation between fibrous cap development and the duration of lipid exposure as well as plaque volume^22,40^. We focussed our analysis on plaques that included lineage-labelled VSMCs (122/150). A cap structure was observed in the majority of lesions containing Confetti^+^ cells, however, a substantial fraction (36%) of lesions with Confetti^+^ cells did not have an obvious cap structure. Plaques lacking a fibrous cap were observed more frequently at the earlier timepoint (42% at W6.5 *vs.* 28% at W11, Figure 1C), also indicating that VSMCs enter the plaque prior to fibrous cap formation.

**Figure 1.**
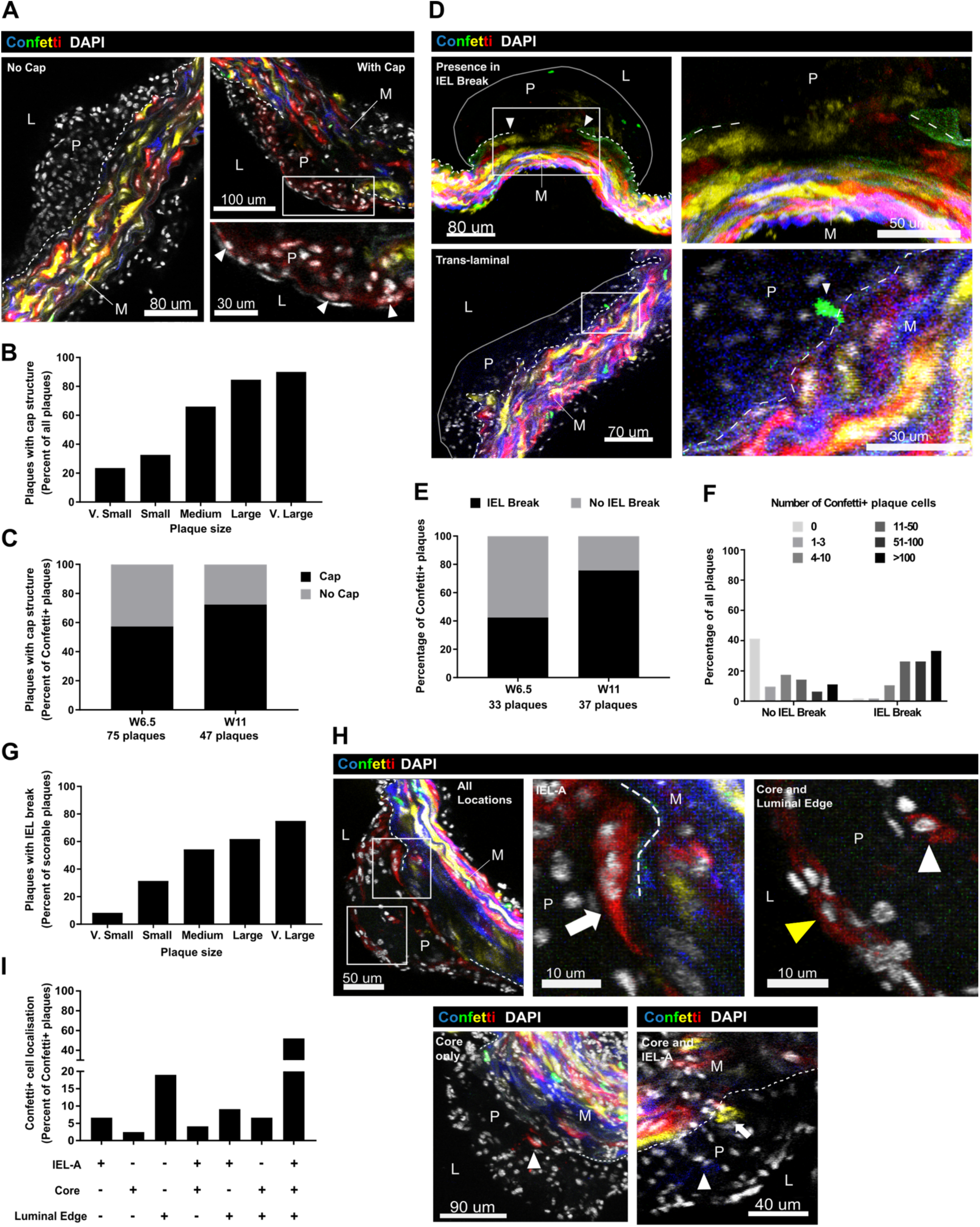
VSMC infiltration in atherosclerosis occurs prior to generation of a fibrous cap. (*A*) Representative confocal images (single *Z*-stack) of plaques without (left) and with fibrous cap structure containing contiguously arranged, elongated DAPI-stained nuclei adjacent to the endothelium (indicated by arrowheads, right) in VSMC-lineage-labelled, high fat diet (HFD) fed Myh11^Confetti^; *Apoe^-/-^* animals. Confetti (CFP: blue, RFP: red, YFP: yellow, GFP: green) and DAPI (white) signals are shown as indicated. The bottom right panel shows a magnified view of the boxed region. Dashed lines show outline of the internal elastic lamina (IEL). Scale bars represent 80 μm (left), 100 μm (top right) and 30 μm (lower right). (*B*) Percent of all plaques with fibrous cap structure according to plaque size. n=150 plaques (n=17 “very (v) small” plaques, 49 “small”, 47 “medium”, 27 “large”, 10 “v. large”). (*C*) Percent of plaques containing Confetti^+^ cells at 6.5 or 11 weeks (W) of HFD feeding. n=122 plaques (n=75 plaques from 4 animals at W6.5 and n=47 plaques from 3 animals at W11). (*D*) Representative confocal images (max. projection) of plaques from HFD-fed Myh11^Confetti^; *Apoe^-/-^* mice, showing Confetti^+^ cells located at a site of IEL break (top panels) and “translaminal” Confetti^+^ cells crossing an intact IEL (lower panels). The right-hand panels show magnified views of the boxed regions. Signals from the Confetti reporter proteins are shown and lower panels also show DAPI signal (white). Solid grey lines show the outlines of plaques and dashed lines show the outline of the IEL. Arrowheads highlight breakpoints of the IEL (top) and a translaminal Confetti^+^ cell (bottom). Scalebars represent 80 μm (top-left), 50 μm (top-right), 70 μm (bottom-left) and 30 μm (bottom-right). (*E*) Proportion of Confetti^+^ cell-containing plaques with an IEL break, stratified by weeks (W) of HFD. n=70 plaques (n=33 plaques from 4 animals at W6.5 and n=37 plaques from 3 animals at W11). (*F)* Distribution of plaques according to the number of Confetti^+^ cells per plaque, for lesions without (54 plaques) or with an IEL break (43 plaques). (*G*) Percent of plaques that have an IEL break, stratified by plaque size. n=9 “v. small” plaques, 29 “small”, 35 “medium”, 17 “large”, 7 “v. large”. Plaques were excluded in (F, *G*) if IEL status could not be confidently scored. (*H*) Representative confocal images (top panels, single Z-plane; lower panels, max. projection) of plaques from HFD-fed Myh11^Confetti^; *Apoe^-/-^* mice, showing Confetti^+^ cells in all plaque regions (top), core only (lower left), or core and “IEL-adjacent” (“IEL-A”)(lower right). Magnified views of the plaque shown in top panels (boxed regions) show IEL-A (middle) and “core and luminal edge” located Confetti^+^ cells (right). Signals for the Confetti reporter proteins and DAPI are shown. White arrows indicate Confetti^+^ cells localising to the IEL-A, yellow arrowhead point to Confetti^+^ cells at the luminal edge and white arrowheads show Confetti^+^ cells in the plaque core. Dashed lines indicate outline of the IEL. (*I*) Distribution of plaque regions invested by Confetti^+^ cells in all plaques with Confetti^+^ cell investment (n=122 plaques). (*A*, *D* and *H*) L, lumen; P, plaque, M, media.

To understand the events preceding generation of the fibrous cap, we examined the localisation and numbers of VSMC-derived cells in developing lesions. First, we considered VSMC entry pathways into plaques and found that in 42 of the 70 Confetti^+^ plaques (60%) where definitive scoring was possible, a break in the internal elastic lamina (IEL) was observed (Figure 1D and E). However, the IEL was intact in 40% of lesions, and no IEL break was the most frequent observation in plaques analysed at W6.5 (60%; Figure 1E). In two cases, we detected a trans-laminal lineage-labeled cell in lesions without IEL breaks (Figure 1D). While this was a relatively rare event, it is in keeping with previous high-resolution confocal microscopy analysis that showed migration of VSMCs through IEL fenestrations^41^. These findings suggest that VSMC plaque infiltration can occur both via cell migration through an intact IEL from the underlying media and through IEL breaks. Plaques without detectable IEL breaks tended to contain lower numbers of VSMC-lineage cells (Figure 1F), indicating that trans-IEL migration is associated with earlier stage plaques with low VSMC investment. On the other hand, plaques with observable IEL breaks contained more VSMC lineage cells (Figure 1F), suggesting entry via this route occurred in more advanced plaques with more VSMC investment, or that IEL breaks are secondary to VSMC investment. This notion is in keeping with an increased frequency of IEL break-containing plaques at W11 (29/42, 69%) compared to W6.5 (14/52, 27%), when considering both Confetti^+^ and Confetti^-^ plaques. Stratification of plaques also showed a plaque size-dependent increase in IEL-break frequency (Figure 1G), further suggesting that IEL breaks are associated with plaque maturity.

To understand how VSMC localisation within plaques changes during lesion development, plaques were scored for the presence of VSMCs in each of three plaque regions: “IEL-adjacent” (“IEL-A”), which include cells in the plaque core that are in direct contact with the IEL, “luminal edge”, which could be part of the fibrous cap, and “core” (Figure 1H). Plaques with VSMCs in all three locations were most common (63 out of 121 scorable Confetti^+^ lesions, 52%), although, we found VSMC investment limited to specific regions in almost half of the plaques (Figure 1I). We observed VSMCs selectively at the luminal edge in 23 plaques (19%, Figure 1I), which is in keeping with a previous study that suggested VSMCs initially migrate through the lesion via the luminal edge^15^. However, a substantial subset of plaques did not have lineage-labelled VSMC contribution at the luminal edge (16 lesions, 13%), including 5 lesions with labelled cells selectively in the region adjacent to the IEL (IEL-A, Figure 1I). Overall, these findings suggests that, while migration via the luminal edge is prevalent, alternative routes exist and demonstrates that VSMCs are present in plaques before cap formation.

### 3.2. Inference of a VSMC transition that gives rise to a fibrous cap-associated cell state in vascular disease models

While the VSMC origin of fibrous cap cells is well documented, the pathways underlying their generation are not fully understood. We therefore examined a scRNA-seq dataset containing lineage-traced VSMCs from atherosclerotic mouse arteries sampled at early, mid and late-stage atherosclerosis^20^ (8, 16 and 22 weeks of HFD on *Apoe^-/-^*background; GSE155513), to enable the capture of active VSMC transitions underlying fibrous cap development. To highlight clusters that represent VSMCs from healthy artery regions, we integrated this dataset with profiles of VSMCs from healthy vessels in control Apoe^+/+^ mice. Athero (A)-cluster 0, 1, 7, 9 and 11 were present in both healthy and atherosclerotic arteries and all expressed typical contractile VSMC (cVSMC) markers (*Myh11, Acta2, Cnn1;* Figure 2A-C; Supplementary Figure 1). As expected, we also detected clusters containing VSMC-lineage cells with downregulated contractile genes in previously described disease-specific states^11,17,19,20,25^. This included an activated, or intermediate modulated VSMC state (imVSMC), characterised by ECM synthesis and expression of *Ly6a, Vcam1, Tnfrsf11b, Lgals3* (A-cluster 4 and 5; Figure 2A-E). These imVSMCs have also been termed “SEM”^19,20^, fibromyocytes^25^ or pioneer cells^21^. Additionally, *Sox9^+^, Col2a1^+^* and *Chad^+^*chondromyocytes (CMCs; A-cluster 2 and 10) and fibroblast-like (FB-like, A-cluster 8) cells expressing high levels of *Pi16, Igfbp4* and *Dpep1*^42^ were detected (Figure 2A and B). The VSMC origin of a distinct cell population expressing *Cd68* and other macrophage markers (MΦ-like; A-cluster 3 and 12; Figure 2 A and B) has been contested^25,42^. We also observed a small population of proliferating (Prlf) VSMCs (*Ccnd1^+^, Pcna^+^* and *Mki67^+^*), which co-expressed imVSMC markers (A-cluster 13, Figure 2A, B and D), in keeping with previous work suggesting that the imVSMC state is associated with proliferation^20^.

**Figure 2.**
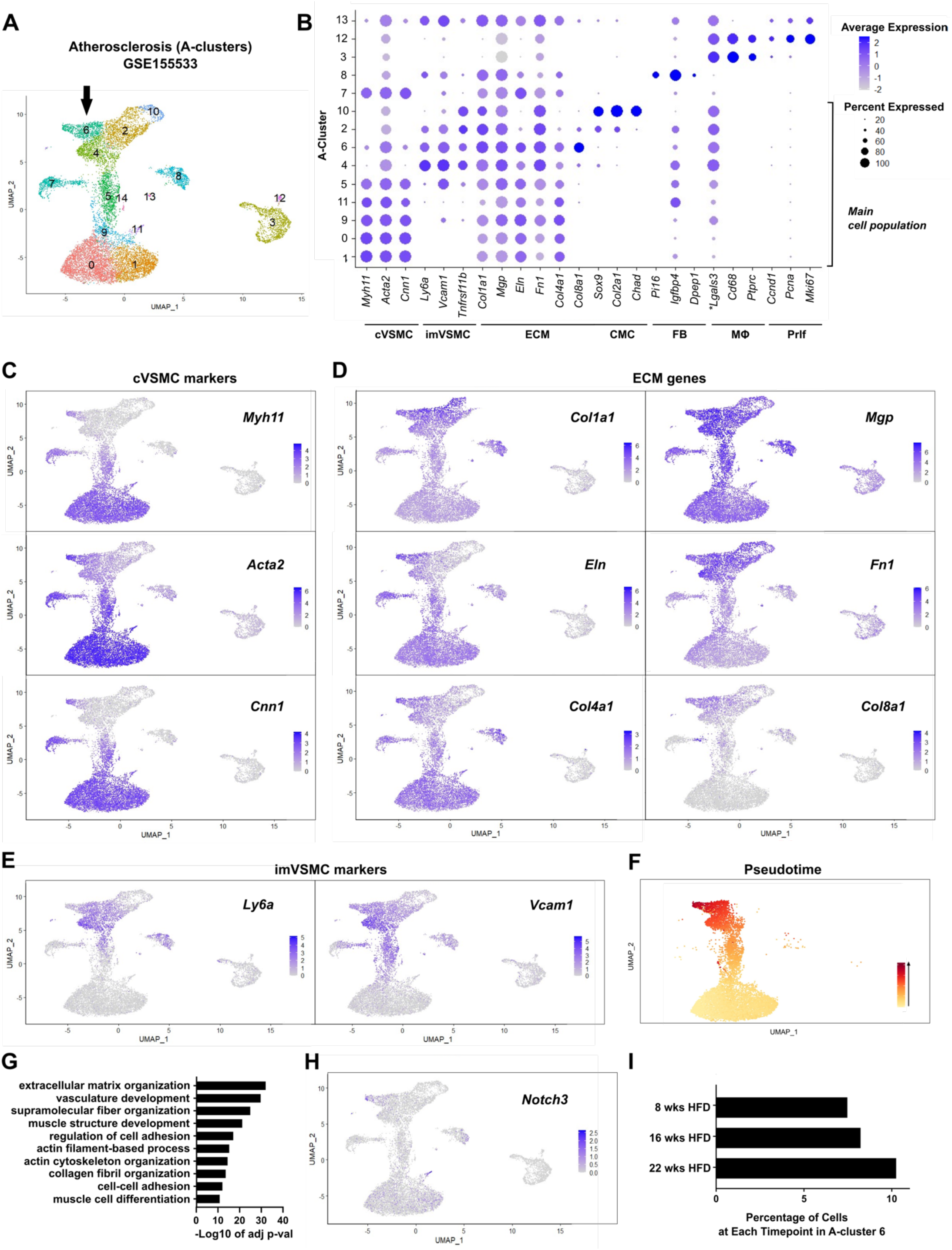
Identification of a proposed fibrous cap-associated VSMC state derived from intermediate modulated VSMCs (*A*) Uniform Manifold Approximation and Projection (UMAP) with atherosclerosis cell cluster (A-cluster) annotation for scRNA-seq analysis of VSMC-lineage label-positive cells from atherosclerotic arteries of Myh11-CreER^t2^; reporter-ZsGreen; *Apoe^-/-^* animals after 8, 16 and 22 weeks of high fat diet (HFD) feeding (GSE155513)^20^ and healthy control Apoe^+/+^ carotid arteries. An arrow points to the proposed fibrous cap-associated cluster, A-cluster 6. (*B*) Dot plot showing expression of cell state markers in clusters using a grey (low) to blue (high) scale and the percentage of cells with detected marker expression for each cluster indicated by dot size. Clusters comprising the main, contiguous cell population are marked. cVSMC: contractile VSMC; imVSMC: intermediate modulated VSMC; ECM: extracellular matrix; CMC: chondromyocyte; FB: fibroblast; MΦ: macrophage; Prlf: proliferation. \**Lgals3* is also expressed by imVSMCs. (*C-E*) UMAP feature plots showing expression levels for cVSMC (*C),* ECM *(D)*, and imVSMC genes (*E)* using grey (low) to blue (high) scales. (*F*) UMAP of only the cells that are part of the Slingshot trajectory resulting in A-cluster 6 cell generation. Pseudotime is indicated using a yellow-red scale. (*G*) Gene ontology (GO) terms enriched for genes upregulated in A-cluster 6 compared to A-cluster 4 cells [P-adj<0.05, log(fold-change)>0.5]. (*H*) UMAP feature plot showing *Notch3* expression using a grey-blue scale. (*I*) Proportion of VSMC-lineage cells in A-cluster 6 for each HFD timepoint.

Interestingly, we detected an additional disease-specific cell cluster (A-cluster 6) that co-expressed contractile VSMC markers (*Myh11, Acta2, Cnn1*) and key ECM genes (*Col1a1, Mgp, Eln, Fn1, Col4a1, Col8a1*; Figure 2A-C and E, Supplementary Figure 1). These cells also expressed *Ly6a, Vcam1 and Tnfrsf11b*, but at a lower level compared to imVSMCs (Figure 2A-E). Consistent with the idea that imVSMCs represent the dedifferentiated VSMCs that migrate into lesions and give rise to the different VSMC-lineage plaque cells^17,18,20,21^, slingshot trajectory analysis predicted that imVSMCs act as an intermediate state for generation of A-cluster 6 cells (Figure 2F). Enrichment analysis of genes that were upregulated in A-cluster 6 cells compared to imVSMCs (A-cluster 4) identified gene ontology (GO) terms associated with contractile VSMC function (“muscle structure development”, “actin cytoskeleton organization”, “muscle cell differentiation”; Figure 2G and Supplementary Table 3), demonstrating an induction of the broader contractile gene program in A-cluster 6. We also detected significant upregulation of genes associated with ECM production (“extracellular matrix organization”, “collagen fibril organization”), suggesting that these cells exhibit a synthetic phenotype. *Notch3,* that has been shown to play an important role in fibrous cap formation^26^, was among the genes showing higher expression in A-cluster 6 than in A-cluster 4 cells (Figure 2H). This expression profile led us to hypothesise that A-cluster 6 cells were associated with fibrous cap formation. Consistent with the association of fibrous cap formation with later disease stages^22,40^ (Figure 1B and C), A-cluster 6 cells were present at a greater proportion at the later timepoint of HFD feeding (22 weeks; Figure 2I and Supplementary Figure 1).

The induction of contractile VSMC markers during the transition from imVSMCs to the proposed fibrous cap-associated VSMC state (referred to as “fcVSMCs”) in atherosclerosis was reminiscent of VSMC “re-differentiation”. We therefore compared this to VSMC regulation in the carotid ligation injury model, which involves reproducible and acute VSMC de-differentiation, formation of a VSMC-rich intimal lesion and subsequent VSMC redifferentiation^6,43,44^. We performed scRNA-seq on VSMC lineage (eYFP^+^) cells sorted from ligated arteries of Myh11-CreERt2/Rosa26-eYFP (Myh11^eYFP^) mice 11 days post injury (DPI) – a timepoint where neointimal lesions are developing and VSMC re-differentiation is ongoing^6,43,44^. Clusters containing VSMC-lineage cell states previously described in injured arteries were detected^18,45^, including *Ly6a^+^*, *Vcam1^+^*, *Tnfrsf11b^+^* and *Lgals3^+^* imVSMCs [injury cell cluster (I-cluster) 3 and 4] and *Ccnd1^+^, Pcna^+^* and *Mki67^+^* Prlf cells (I-cluster 9; Figure 3A-E). Interestingly, we also detected cells with high levels of *Sox9, Col2a1* and *Chad*, similar to CMCs (I-cluster 15). Rare *Pi16^+^, Igfbp4^+^*and *Dpep1^+^* FB-like (I-cluster 12) and *Lgals3^+^, Cd68^+^* and *Ptprc^+^* MΦ-like cells (I-cluster 14) showed low expression of the lineage label, *eYFP* (Figure 3A-E), suggesting that these cells were not derived from VSMCs, similar to what has been proposed in atherosclerosis^42^. Cells with high *Myh11, Acta2* and *Cnn1* expression were classed as cVSMCs (I-cluster 0, 1, 2, 7, 10, 11).

**Figure 3.**
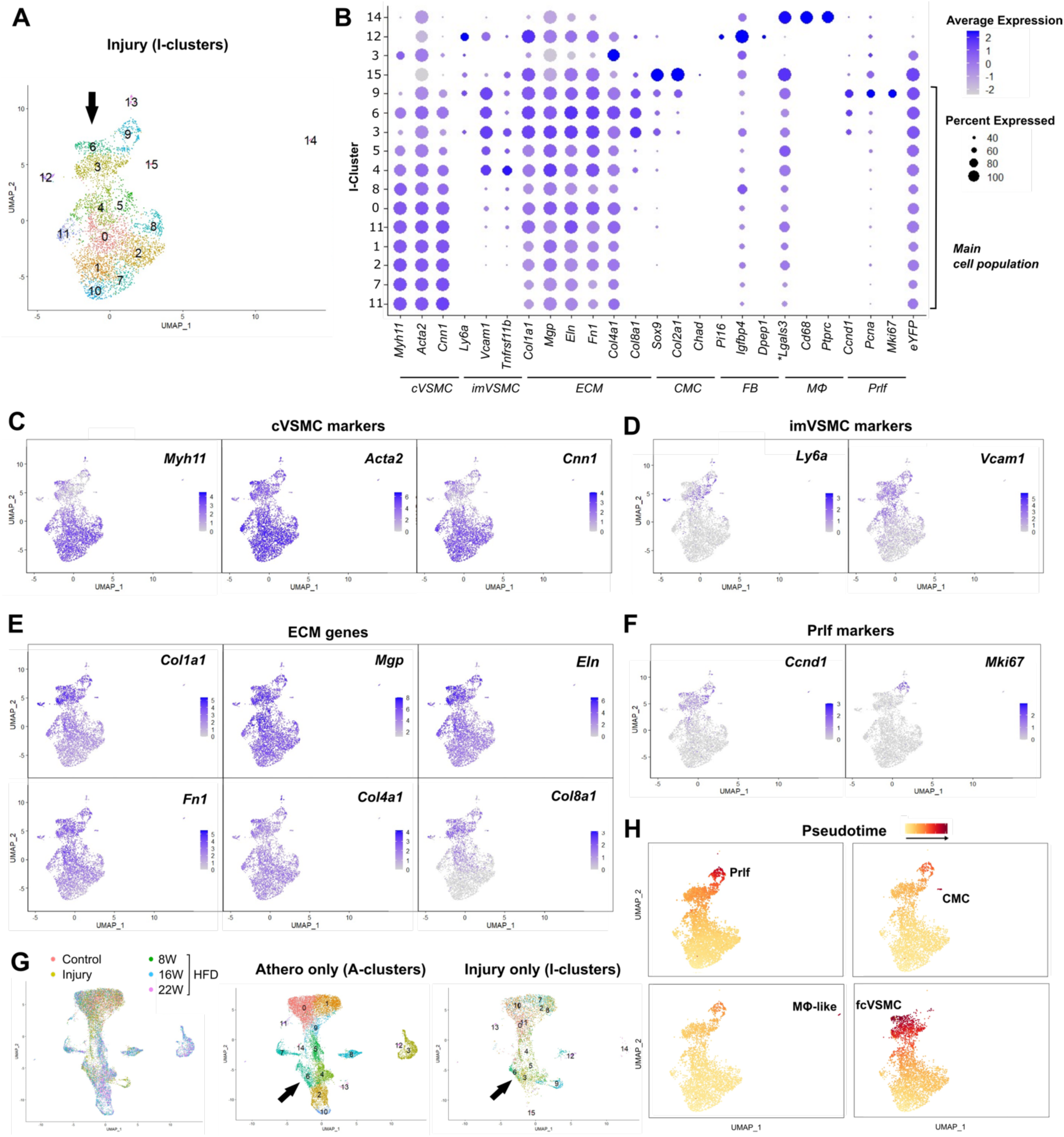
VSMC differentiation from an intermediate modulated state after carotid ligation injury also generates the transcriptional fibrous cap-associated VSMC state. (*A*) Uniform Manifold Approximation and Projection (UMAP) showing injury (I) cell cluster annotation for scRNA-seq analysis of VSMC-lineage label positive cells from ligated left carotid arteries of Myh11^eYFP^ animals 11 days post injury (DPI). An arrow points to the proposed fibrous cap-associated cluster (I-cluster 6). (*B*) Dot plot showing expression of cell state markers and the eYFP lineage reporter in clusters using a scale from grey (low) to blue (high) and the percentage of cluster cells with detected marker expression indicated by dot size. Clusters comprising the main, contiguous cell population are marked. cVSMC: contractile VSMC; imVSMC: intermediate modulated VSMC; ECM: extracellular matrix; CMC: chondromyocyte; FB: fibroblast; MΦ: macrophage; Prlf: proliferation. \**Lgals3* is also expressed by imVSMCs. (*C-F*) UMAP feature plots showing expression levels for markers of cVSMC (*C),* imVSMC (*D*), ECM (*E)*, and proliferation *(F)* using grey (low) to blue scales (high). (*G*) Integration of VSMC scRNA-seq profiles from atherosclerosis [8, 16, 22 weeks high fat diet (HFD), GSE155513^20^], healthy control carotid arteries and the injury (11 DPI) datasets. Left panel shows all cells coloured by dataset identity; middle panel shows cells from the atherosclerosis data only coloured by A-clusters (see Figure 2); right panel shows cells from the 11 DPI data only coloured by I-clusters. *(H)* UMAP showing cells that are part of the Slingshot trajectories leading to generation of a Prlf (I-cluster 9), CMC (I-cluster 15), MΦ-like or fibrous cap-associated (fcVSMC; I-cluster 6) cell state. Pseudotime is indicated using a yellow-red scale.

We also identified a cell population co-expressing contractile genes with key ECM genes (*Col1a1, Mgp, Eln, Fn1, Col4a1, Col8a1*) and *Notch3*, similar to the fcVSMC population in the atherosclerosis dataset (I-cluster 6; Figure 3A-E, Supplementary Figure 2). This population was not detected at earlier timepoints following injury (5 and 7 DPI)^18^, suggesting that these cells are derived from modulated cells and represent VSMC re-differentiation. Integration of the datasets mapped I-cluster 6 to the fcVSMC population in the atherosclerosis data (A-cluster 6; Figure 3F). Like in atherosclerosis, computational trajectory analysis indicated that I-cluster 6 cells were derived from imVSMC cells, which was confirmed using partition-based graph abstraction (PAGA) as an alternative inference method (Figure 3H and Supplementary Figure 3). Cells from the two disease models mapped together after integration of the data (Figure 3G, H), suggesting that the range of VSMC states induced by carotid ligation injury recapitulates those observed in atherosclerosis. Interestingly, the trajectory inference analysis suggested that imVSMCs act as an intermediate state for other VSMC-derived subsets in injury (Figure 3H), similar to what has been proposed in atherosclerosis^19,20^. An expected transition to the Prlf state via imVSMC state was also present (Figure 3G). Together, these data support the idea that VSMCs transition to disease-relevant states via an imVSMC state and suggest that these transitions can be investigated using the reproducible vascular injury model. Importantly, our analysis identified a novel imVSMC-derived fcVSMC state in both models, that we hypothesise is implicated in fibrous cap formation.

### 3.3. VSMC transition to a NOTCH3^+^ fibrous cap-associated cell state occurs in the fibrous cap and injury-induced lesions

To assess whether the imVSMC-to-fcVSMC transition captures generation of the fibrous cap, we developed a strategy to co-stain Confetti^+^ samples for NOTCH3 and VCAM1. We used NOTCH3 to mark the fcVSMC state, as *Notch3* expression was largely confined to fcVSMCs in both atherosclerosis and injury (Figure 2H and Supplementary Figure 2). *Vcam1* was highly expressed in imVSMCs and showed downregulation in fcVSMCs (Figure 2B and D), hence, VCAM1^+^ NOTCH3^-^ cells highlighted the imVSMC state. VCAM1^+^NOTCH3^+^ cells represent cells at the cluster boundary (A-cluster 4-to-6 or I-cluster 3-to-6). Clonally related VSMC-derived cells express the same fluorescent reporter protein in Myh11^Confetti^ mice, hence, this approach allowed us to assess the lineage relationship of cells present in these states. We scored 26 individual patches of VSMC-derived cells of the same colour in 16 plaques from three Myh11^Confetti^; *Apoe^-/-^* animals for the presence of cells expressing each marker combination (Figure 4A-C). VSMC clones containing cells with all marker combinations (VCAM1^+^NOTCH3^-^, VCAM1^+^NOTCH3^+^ and VCAM1^-^NOTCH3^+^ cells) were most frequently observed (42.3%; Figure 4B), showing that the progeny of a single VSMC can generate all cell states involved in the imVSMC-to-fcVSMC transition. We did not observe VSMC clones containing only VCAM1^-^NOTCH3^+^ cells in any plaques, whilst VCAM1 was detected without NOTCH3 in 31% of clones (Figure 4B). This is consistent with the idea that VCAM1 induction is associated with VSMC entry into the plaque and that cells subsequently transition to the NOTCH3^+^ state. We hypothesised that the local plaque environment would impact the likelihood of generating the imVSMC and fcVSMC states such that cells in the fibrous cap would have a greater propensity to adopt the fcVSMC state. To test this idea, we quantified marker-expressing cells in each VSMC clone separately for the luminal region (including clones exclusively located at the luminal edge and luminally-located cells in clones spanning both core and cap) and the core region (clones exclusively located in core and core-located cells in clones spanning both core and cap). In keeping with previous observations^26^, we found abundant NOTCH3 expression by VSMCs in the luminal region, whereas VCAM1 expression was more prominent in the core region of VSMC clones (Figure 4A and C). Similar proportions of the three cell states were detected in the luminal region, suggesting this is the site of imVSMC-to-fcVSMC state transition. Together, these data shows that the computationally inferred transition from imVSMC to fcVSMC state occurs *in vivo*, and that this represents formation of the fibrous cap in atherosclerotic plaques.

**Figure 4.**
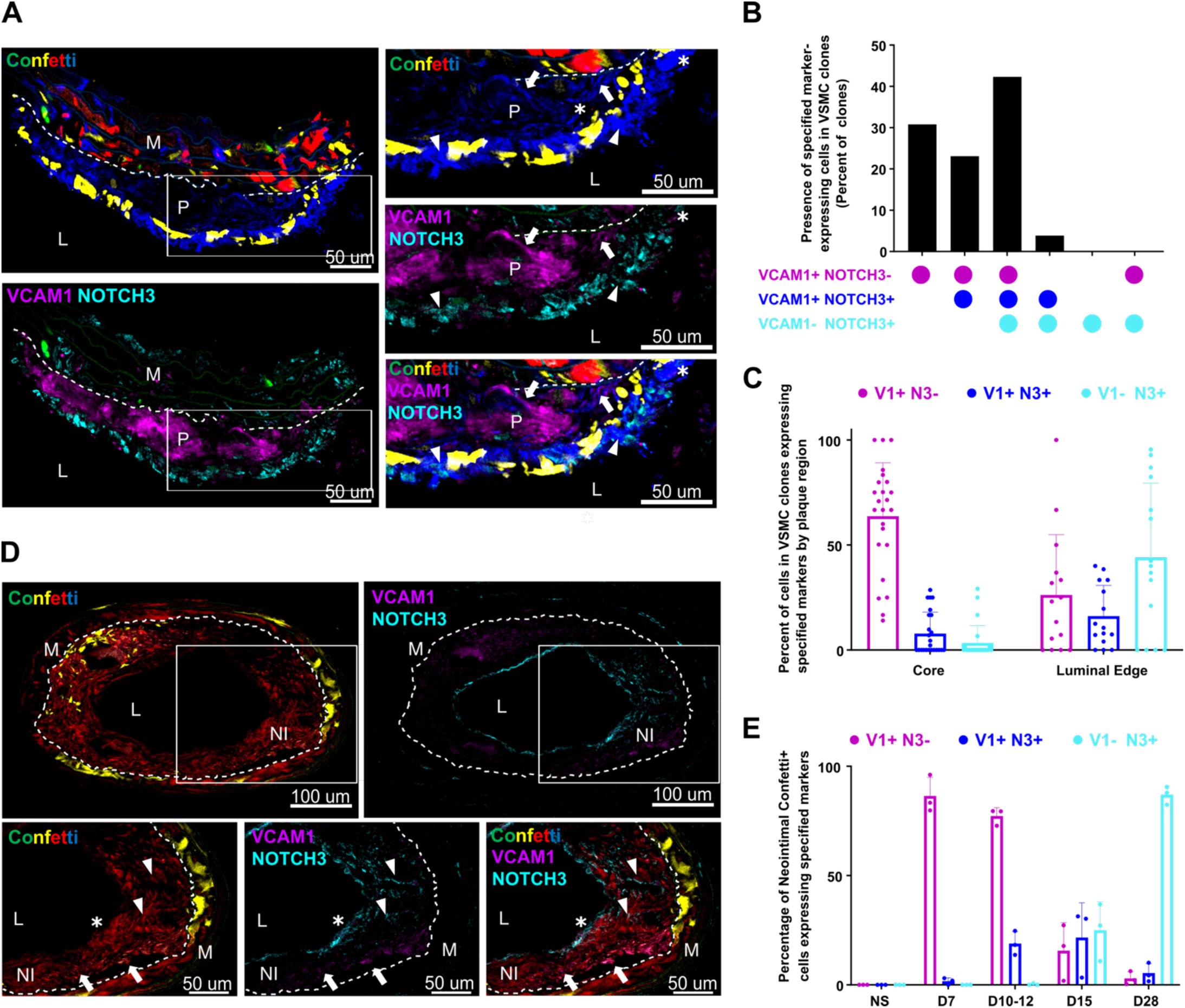
VSMCs in the intermediate modulated and fibrous cap-associated state co-occur in clonally related cells within lesions. (A-C) Immunofluorescence staining for VCAM1 and NOTCH3 in cryosections of a plaque from VSMC-lineage-labelled Myh11^Confetti^; *Apoe^-/-^*animals. *(A)* Representative image of plaque. Panels show the entire plaque section (left) and magnified views of boxed region (right). Signals for VCAM1 (magenta), NOTCH3 (cyan) and Confetti (CFP: blue, RFP: red, YFP: yellow, GFP: green) are shown as indicated. Examples of VCAM1^+^ (arrows), NOTCH3^+^ (arrowheads) and VCAM1^+^NOTCH3^+^ cells (asterisk) are indicated. Dashed lines outline the internal elastic lamina (IEL). Scale bars = 50 µm. L, lumen; P, plaque, M, media. (*B*) Percentage of plaque-located Confetti^+^ clones that contain cells with specified combinations of markers expressed. n=26 clones from 3 animals. (*C*) Percentage of cells with specified combinations of marker expression quantified separately for either the core or luminal edge portion of individual Confetti^+^ clones. n=23 (core region) and 15 clones (luminal edge). Bars show mean, error-bars SD. (*D-E*) Immunofluorescence staining for VCAM1 and NOTCH3 in cryosections of the ligated artery of VSMC-lineage-labelled Myh11^Confetti^ animals. (*D*) Representative image at 15 days post injury. Panels show the entire section (top; scale bars = 100 µm) and magnified views of boxed regions (bottom; scale bars = 50 µm). Signals for VCAM1 (magenta), NOTCH3 (cyan) and Confetti (CFP: blue, RFP: red, YFP: yellow, GFP: green) are shown as indicated. Arrows point to VCAM1^+^ cells in the neointima, arrowheads point to NOTCH3^+^ cells in the neointima and an asterisk marks an example of a cell expressing both VCAM1 and NOTCH3. Dashed lines outline the IEL. L, lumen; NI, neointima, M, media. (*E*) The proportion of neointimal Confetti^+^ cells with specified marker expression at different timepoints after injury (n=3 animals per timepoint). Bars show mean, error-bars SD.

To more directly test whether lesional NOTCH3^+^ fcVSMC cells are generated from VCAM1^+^ imVSMCs, we used the carotid ligation injury model, where the acute and reproducible kinetics of lesion formation could be leveraged. Similar to the analysis of plaques, VSMC clones containing VCAM1^+^NOTCH3^-^, VCAM1^+^NOTCH3^+^ and VCAM1^-^NOTCH3^+^ cells were also detected in injury-induced lesions (Figure 4D), however, the frequency of VSMC state marker profile varied over time. At initial stages of lesion formation (7 DPI), almost all lesional VSMCs (86%) were VCAM1^+^NOTCH3^-^(Figure 4E). VCAM1 was also expressed by VSMCs in the media of diseased artery regions at 7 DPI; in contrast, NOTCH3 expression was detected in very few cells at this timepoint. A small reduction in intimal VCAM1^+^NOTCH3^-^ cells (to 77%), coupled with a substantial proportion of VCAM1^+^ NOTCH3^+^ cells was observed at D10-12 (19%). At 15 DPI, VCAM1^+^NOTCH3^-^, VCAM1^+^NOTCH3^+^ and VCAM1^-^NOTCH3^+^ states were detected at similar frequency (∼20% each; Figure 4E). This overlaps the reported re-expression of contractile genes at 2 weeks after injury in this model^44^. In contrast, almost all cells were VCAM1^-^NOTCH3^+^ at 28 DPI (87%; Figure 4E). Analysis of cells in the medial layer of injured arteries showed NOTCH3 re-expression by VSMCs at 15 and 28 DPI, consistent with VSMC redifferentiation of medial VSMCs (Supplementary Figure 4). Interestingly, we did not detect VCAM1^+^NOTCH3^+^ cells in the medial layer at any of these timepoints (Supplementary Figure 4), suggesting that the kinetics or underlying mechanisms may differ. Together, these data demonstrate that VSMCs enter the lesion in an imVSMC state and that clonal progeny of individual VSMCs transition to the fcVSMC state within injury-induced lesions.

### 3.4. The thrombin receptor PAR1 is a candidate therapeutic target for promoting the fibrous cap-associated VSMC transition

Next, we aimed to understand the regulatory mechanisms governing the generation of the fcVSMC state, which could be explored for promoting fibrous cap formation and thus plaque stabilisation. To this end, we found 840 genes showing significant changes in expression (*P*-adj<0.05, log[fold-change]>0.25) along pseudotime for the trajectory generating fcVSMCs in atherosclerosis (Figure 5A). These overlapped significantly with genes showing pseudotime-associated expression for the trajectory involving imVSMC to fcVSMC transition after injury (59%; Chi-squared test, p<0.001; Figure 5B), further demonstrating the similarity of VSMC regulation in these models. Hierarchical clustering of pseudotime-associated genes by expression pattern highlighted a gene subset (A-clade 3) specifically upregulated at the boundary of A-cluster 4 and 6 cells, representing the imVSMC to fcVSMC transition (boxed in Figure 5A). The majority (67%) of these 83 imVSMC-to-fcVSMC transition-induced genes were similarly upregulated along the pseudotime of the corresponding trajectory in injury (I-clade 2; Figure 5A and B). We focused on these 56 genes with similar expression profile in the injury model, hypothesising that the acute, accelerated kinetics of lesion formation in this model would enrich for genes that drive fcVSMC differentiation, rather than being expressed as a consequence of the transition.

**Figure 5.**
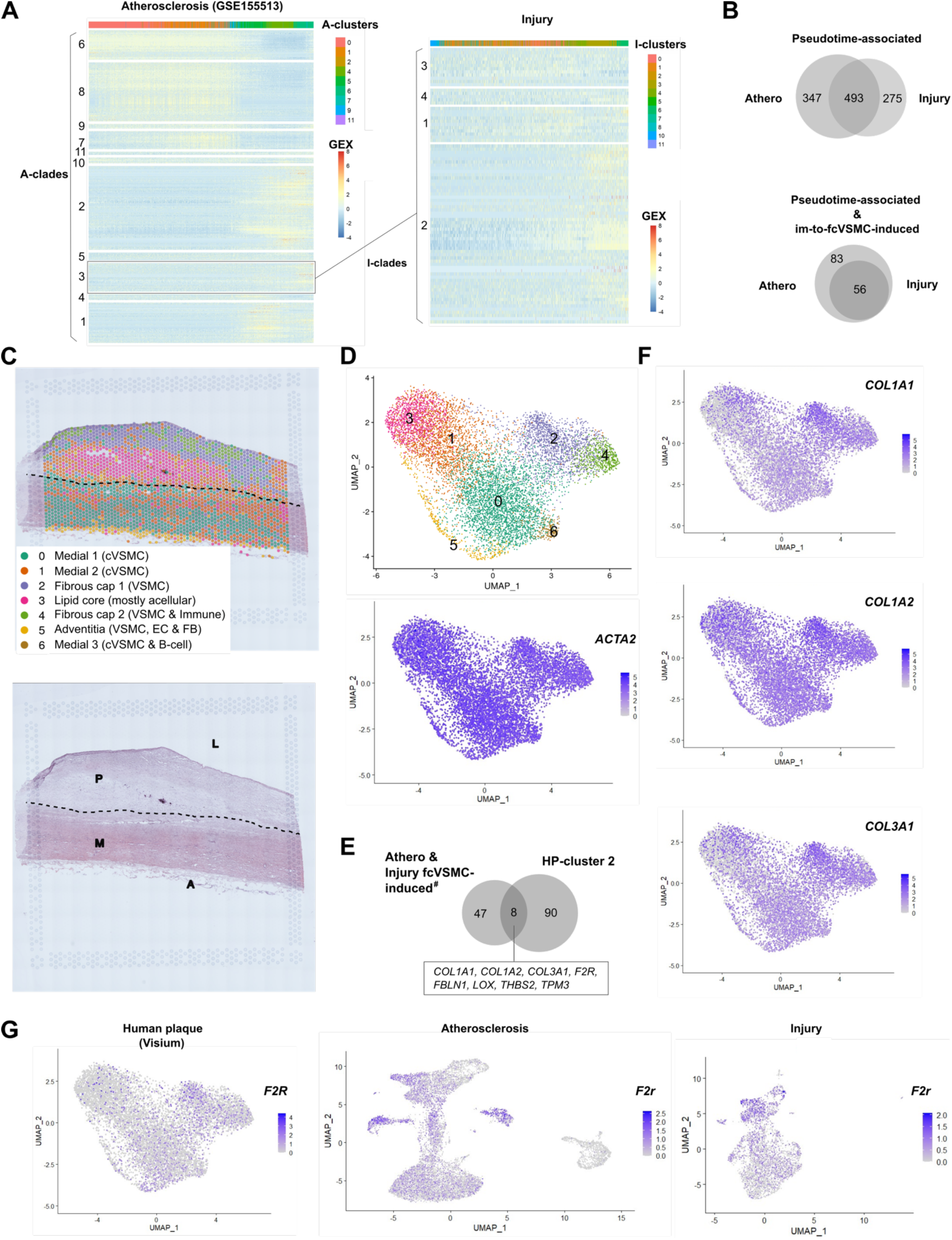
Identification of PAR1 as a candidate regulator of VSMC differentiation to a fibrous cap-associated VSMC state. (*A*) Left panel shows heatmap of genes with significant pseudotime-associated expression for the trajectory generating the fibrous cap-associated (fcVSMC) A-cluster 6 [P-adj<0.05, log(fold-change)>0.25], clustered into 11 “A-gene clades” based on correlated gene expression patterns. The boxed gene clade (A-3) is upregulated at the A-clusters 4-6 border. Right panel shows the expression of A-clade 3 genes along the fcVSMC-generating trajectory in the injury data (I-cluster 6) after re-clustering according to expression in the injury dataset (I-gene clades). Top bars show cell cluster affiliation using colour scale from Figure 2A and 3A for the atherosclerosis and injury datasets, respectively. Scaled gene expression is shown by a colour scale from blue (low) to red (high). (*B*) Venn diagrams showing the overlap of all pseudotime associated genes for the fcVSMC paths in the atherosclerosis and injury datasets (top) and the overlap of fcVSMC-induced A-clade 3 and I-clade 2. (*C*) Spatial transcriptomics analysis of human plaque from a 54-year-old male organ donor, showing the haematoxylin and eosin-stained section overlayed by capture spot cluster annotations [human plaque (HP)-cluster 0-6, top] or labelled by tissue regions (bottom). L, lumen; P, plaque, M, media; A, adventitia. (*D*) Uniform Manifold Approximation and Projection (UMAP) of capture spots coloured by HP-cluster affiliation (top) and *ACTA2* expression(bottom). (*E*) Venn diagram showing the overlap of the 56 fcVSMC-induced genes in both atherosclerosis and injury (I-clade 2) and fibrous cap-located HP-cluster 2. The 8 overlapping genes are listed. (*F*) UMAP feature plots showing fibrillar collagen gene expression. (*G*) UMAP feature plots of *F2R* gene expression in the human plaque (Visium; left), mouse atherosclerosis (middle) and mouse vascular injury datasets (right). (*D*, *F* and *G*) Feature plots show gene expression levels using a grey (low) to blue (high) colour scale.

To assess the relevance of these genes to fibrous cap formation in human atherosclerosis, we performed spatial transcriptomics analysis in an aortic sample with an early-stage, fibrous cap-containing plaque (Figure 5C). This ensured that ongoing fibrous cap formation was likely, thereby enabling the elucidation of active gene regulatory mechanisms driving this process. Capture spots formed 7 clusters (human plaque [HP]-cluster 0-6) that showed regional specificity and that were all indicated to have some VSMC contribution based on *ACTA2* expression (Figure 5C and D). HP-cluster 0 and 1 were located in the media and expressed contractile VSMC markers at high levels (*MYH11, ACTA2, CNN1;* Supplementary Figure 5). Low-abundance HP-cluster 6 spots in the media expressed immunoglobulin genes (*IGHG1, IGKC, IGLC2*) and contractile VSMC markers, indicating VSMC and B cell presence. HP-cluster 5 spots localised to the adventitia and expressed a mix of VSMC, endothelial (*PECAM1, VWF*) and fibroblast (*DCN, FBLN1, IGFBP4*) markers. HP-cluster 3 spots were characterised by poor QC parameters and predominantly resided within the lipid core, a largely acellular region. HP-cluster 2 and 4 were located in the fibrous cap and expressed markers of modulated VSMCs (*SPP1, LUM, TIMP1, FBLN2, SERPINE1, TNFRSF11B)*. Immune cell markers were also abundantly expressed in HP-cluster 4 (*CD68, CD163, CD86*), but these were detected at lower levels in HP-cluster 2 (Supplementary Figure 5). Thus, we focused on HP-cluster 2, since it was the fibrous cap cluster that contained mainly VSMCs. We found a significant overlap between the 56 fcVSMC-induced genes and genes showing significant upregulation in HP-cluster 2 (90 genes; Chi-squared test, p<0.001). The 8 genes with induced expression in both fcVSMC subsets and the fibrous cap included 3 fibrillar collagen genes associated with fibrous cap formation^46^ (*COL1A1, COL1A2, COL3A1), FBLN1, LOX, TPM3, THBS2 and F2R*; Figure 5E and F; Supplementary Figure 5).

We focused on *F2R*, which encodes the thrombin receptor or protease-activated receptor 1 (PAR1), which showed increased levels in fibrous cap-located HP-cluster 2 (Figure 5G). Fibrous cap association in human atherosclerosis was confirmed by immunostaining, which detected abundant PAR1-expressing ACTA2^+^ VSMCs in human plaque fibrous caps, and PAR1 protein was also detected at lower levels in the core (Figure 6A). *F2r* transcripts were detected at high levels in fcVSMC clusters in both atherosclerosis and injury models, with low expression in A-cluster 7 (previously termed minor VSMC population^20^), fibroblast-like A-cluster 8 cells, and imVSMCs (I-cluster 3; Figure 5G). Previous investigations have shown that thrombin promotes VSMC proliferation^47,48^, but a role in VSMC re-differentiation and fibrous cap generation has not been investigated. We therefore assessed whether PAR1 functionally affects the imVSMC-to-fcVSMC transition and thus potentially fibrous cap formation. To this end, we tested the effect of PAR1 stimulation on contractile gene expression in primary human VSMC isolates following thrombin treatment. We observed a significant and dose-dependent increase in *CNN1* and *ACTA2* expression in response to physiologically relevant thrombin concentrations (Figure 6B). Interestingly, this effect was more profound in VSMCs isolated from female compared to male donors (Figure 6B).

**Figure 6.**
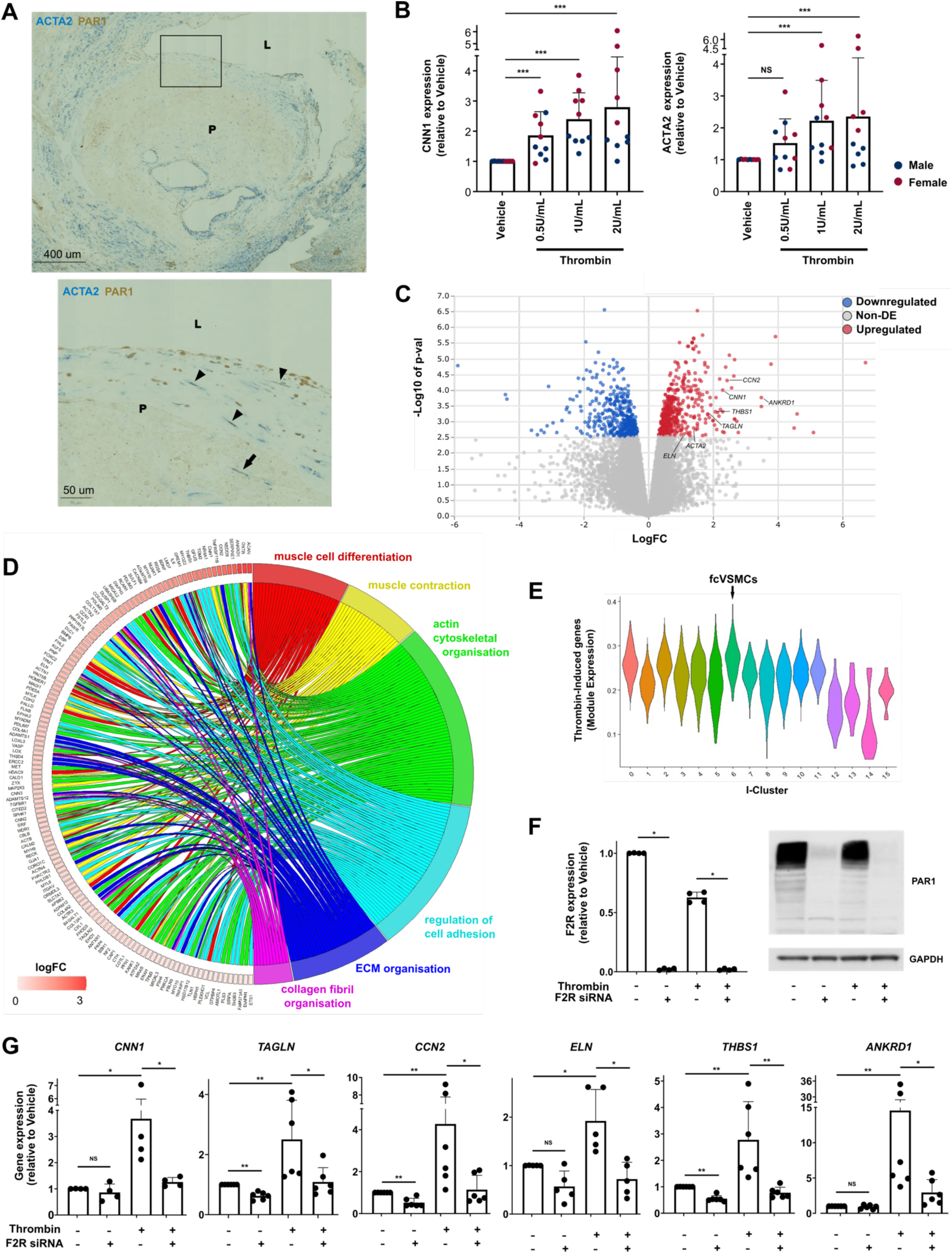
PAR1 is expressed in fibrous cap VSMCs and stimulation of the receptor induces the fibrous cap-associated cell transcriptional pattern. (*A*) Representative immunohistochemistry staining for ACTA2 and PAR1 in sections of human carotid plaques (n=5). Panels show a large region of the plaque (top) and a magnified view of the boxed region (bottom). Signals for ACTA2 (blue) and PAR1 (brown) are shown as indicated. Arrowheads point to examples of ACTA2^+^PAR1^+^ cells in the cap and arrow points to an ACTA2^+^PAR1^+^ cell in the core. Scale bars = 400 µm (top) and 50 µm (bottom). L, lumen; P, plaque, M, media. (*B*) Real-time quantitative reverse transcription PCR (RT-qPCR) analysis of contractile gene expression (*CNN1* and *ACTA2*) in primary human VSMC (hVSMC) isolates from different donors (n=5 male and 5 female donors) after 24 hour vehicle or thrombin treatment (0.5, 1 or 2 U/mL). Male and female donor cell lines are highlighted in blue and red, respectively. Bars show mean ± SD relative to vehicle-treated samples. Statistical testing was done relative to vehicle-treated samples, ***P-adj<0.001; NS: not significant (Mann-Whitney with Bonferroni multiple testing correction). (*C*) Volcano plot of differentially expressed (DE) genes between 24 hour thrombin treatment (2U/mL) and vehicle controls in 3 female donor hVSMC lines (P-adj<0.05) as measured by RNA-sequencing. Red indicates genes upregulated by thrombin treatment, blue indicates downregulated genes and grey indicates non-significant. Genes of interest are indicated. (*D*) Circos plot showing VSMC- and ECM-associated gene ontology (GO) terms (and associated genes) enriched by the 492 genes upregulated by thrombin treatment (from *C*). Differential expression [log(fold-change)] is shown with a white (low) to red (higher in thrombin-treated samples) colour scale. *(E*) Violin plot showing expression of thrombin-induced gene orthologues across the injury dataset cell clusters. The arrow marks the fibrous cap-associated (fcVSMC) I-cluster 6. (*F*) Validation of siRNA-induced *F2R/*PAR1 knockdown in primary human VSMCs in the absence or presence of thrombin. Left panel shows transcript levels normalised to vehicle. Bars show mean ± SD. n=4 hVSMC lines from different donors. *P-adj<0.05 (Mann-Whitney with Bonferroni multiple testing correction). Statistical testing was done between non-targeting control and *F2R/*PAR1 siRNA treated samples). Right panel shows protein levels by representative western blotting for PAR1 (top) and GAPDH (bottom). n=3 hVSMC lines from different donors. (*G*) Expression levels of selected genes of interest in thrombin-treated hVSMCs relative to vehicle-treated samples with and without F2R siRNA knockdown. Bars show mean ± SD. n=4-6 cell lines. *P-adj<0.05, **P-adj<0.01, ***P-adj<0.001, NS: not significant (Mann-Whitney with Bonferroni multiple testing correction, Statistical testing was done between groups).

To more comprehensively evaluate the transcriptional state induced by thrombin, we profiled gene expression in thrombin-treated and untreated cells using RNA-seq. This analysis was performed using cells from 3 female donors, due to the enhanced response observed in female VSMCs. We confirmed upregulation of contractile VSMC genes (*CNN1, ACTA2, TAGLN*) in thrombin-treated *vs.* control cells, and also detected increased levels of genes that have been associated with VSMC differentiation and/or fibrous cap development (Figure 6C). For example, connective tissue growth factor (CCN2) is involved in maintenance of the contractile VSMC state and is expressed in the fibrous cap^49,50^; Elastin (ELN) is a key ECM component of the fibrous cap and also functions to maintain VSMC contractility^40,51^; Thrombospondin 1 (THBS1) has been implicated as a key factor involved in fibrous cap formation^52^; and Cardiac ankyrin repeat protein (ANKRD1) is expressed by VSMCs of the fibrous cap^53^. Systematic analysis showed that the 492 thrombin-upregulated genes (P-adj<0.05) were enriched for GO terms associated with muscle cell differentiation, contraction and focal adhesion, suggesting an overall induction of VSMC differentiation and function-associated genes (Figure 6C and D; Supplementary Table 4). Thrombin-upregulated genes concomitantly showed enrichment for GO terms associated with ECM organisation, including collagen fibril organisation (Figure 6C and D, Supplementary Table 4), which is a key component of fibrous cap formation^46^. Remarkably, a gene module comprising murine orthologues of the genes upregulated by thrombin in human VSMCs exhibited peak expression in the fcVSMC population identified in injured mouse arteries (I-cluster 6; Figure 6E). This suggests an overall induction of a fibrous cap-associated cell state by thrombin treatment of human VSMCs.

To confirm that thrombin-induced expression of contractile and fibrous cap-relevant genes was mediated by thrombin receptor stimulation, we targeted PAR1 activity using siRNA and observed efficient knockdown (Figure 6F). Thrombin-induced expression of *CNN1, TAGLN, CCN2, ELN, THBS1* and *ANKRD1* was inhibited by PAR1 knockdown (Figure 6G). In parallel, we used a PAR1- specific small molecular inhibitor, vorapaxar, which also blocked thrombin-mediated gene induction (Supplementary Figure 6). Interestingly, baseline expression of *TAGLN*, *CCN2* and *THBS1* was reduced by PAR1 knockdown, suggesting thrombin-independent regulation of these genes by PAR1 (Figure 6G). Together, these data indicate that stimulation of the thrombin receptor in VSMCs leads to induction of genes associated with the fcVSMC state. We suggest that, by promoting an imVSMC-to-fcVSMC transition, PAR1 stimulation in VSMCs contributes to fibrous cap formation.

## 4. Discussion

Here, we provide evidence that a novel VSMC sub-population (that we term fcVSMCs) is derived from VSMCs expressing a previously described intermediate modulated, or imVSMC, state, is present in the fibrous cap of plaques and can be modelled in injury-induced lesions. This is supported by I) the demonstration that the fcVSMC transcriptional profile in both atherosclerosis and injury includes key fibrous cap-associated genes, II) trajectory analysis consistently predicts that fcVSMCs are derived from imVSMCs, III) the demonstration that marker combinations representing imVSMCs and fcVSMCs are co-detected in clonally related cells in atherosclerotic lesions, IV) that NOTCH3-expressing intimal cells are not detected until re-differentiation occurs after injury, and V) the fcVSMC marker, NOTCH3, shows abundant expression in the fibrous cap. Based on these findings, we identify candidate regulators of the intermediate-to-fcVSMC transition, including *F2R*- encoded PAR1. We provide evidence suggesting that PAR1 stimulates fibrous cap formation, which includes expression of PAR1 by fcVSMCs and detection of PAR1^+^ VSMCs in the fibrous cap of human lesions. Finally, we show functionally that PAR1 stimulation by thrombin in primary human VSMCs induces fibrous cap VSMC markers and a transcriptional profile matching that of fcVSMCs.

The notion that an imVSMC state gives rise to fibrous cap VSMCs is in line with previous studies. In particular, we previously found that SCA1-expressing VSMCs are present prior to disease and have increased proliferative capacity^18^, similar to the SCA1^+^ cells isolated from lesions^20^. It is therefore conceivable that cells in an imVSMC state invade lesions and expand clonally to generate all VSMC-derived plaque cells, including those comprising the fibrous cap^12^. Other groups have shown that manipulation of factors associated with promoting a VSMC state resembling the intermediate state (downregulation of contractile markers, upregulation of migration- and proliferation-associated genes) disrupt fibrous cap formation^24,25^. Our findings are also consistent with a study using a dual lineage tracing system to monitor VSMC transition through an *Lgals3*-expressing state, a gene which also marks the imVSMC state. Here, all lesional VSMCs, including those in the fibrous cap, were indicated to have transitioned through the intermediate, *Lgals3-* expressing state, at least in early atherogenesis^21^. Notably however, VSMCs that were not of the *Lgals3-*lineage were present in later stage plaques. Martos-Rodriguez et al. found that while NOTCH signalling is lost in VSMCs entering plaques, its reactivation is required for fibrous cap formation^26^. NOTCH signalling is known to promote the contractile VSMC phenotype^54,55^ and thus likely inhibits transition to the intermediate state, which lacks contractile marker expression. This is consistent with our findings, that show upregulation of *Notch3* in fcVSMCs, confirmation of NOTCH3 expression in fibrous cap VSMCs and delayed NOTCH3 expression when compared to VCAM1.

Our finding that PAR1 stimulation induces expression of contractile genes in cultured VSMCs is consistent with data from a published microarray study in human aortic VSMCs after shorter thrombin treatment^56^, although this was not highlighted by the authors. Furthermore, previous work has shown that CCN2, which we found to be upregulated by PAR1 activation, was also induced by thrombin in rat VSMCs^57^. The role of thrombin/PAR1 in VSMCs and vascular disease has been extensively studied^47,48,58^. PAR1 is known to be expressed by VSMCs and thrombin is a well-known mitogen of these cells *in vitro*, putatively via PAR1^47,48^. This proliferation-stimulating effect has been discussed for its potential to exacerbate the response to vascular injury^58^. VSMC proliferation has been detected in the fibrous cap^15^ and it is generally considered to indicate plaque stability, however, the role of PAR1 in fibrous cap formation has not been studied. PAR1 is also expressed by endothelial cells and systemic inhibition of either thrombin or PAR1 attenuated atherosclerosis progression and even promoted features of plaque stability, due to effect on endothelial function, proinflammatory signalling and immune cell infiltration^58-62^. Thus, further investigation of the VSMC-specific effects of thrombin signalling in atherosclerosis is needed, in particular whether this promotes fibrous cap formation in lesions. Indeed, thrombin and several other proteases capable of activating PAR1 (e.g. matrix metalloproteases) are abundant in atherosclerotic plaques^58,59,63,64^. The identification of PAR1 as a candidate regulator of fibrous cap formation could have several implications for the management of atherosclerosis, as several anticoagulants used to treat patients with atherosclerotic disease inhibit either thrombin directly or its receptor, PAR1^65,66^. Given the fibrous cap-inducing effect of VSMC-expressed PAR1 suggested here, such treatment could therefore have an unintended negative effect on plaque stability. Notably, the induction of fcVSMC-associated genes by thrombin was more prominent in female than male VSMCs, and could contribute to the sex differences in plaque biology, particularly, the greater frequency of thin or ruptured fibrous caps in male patients^67,68^.

Our observation that the spectrum of VSMC states observed in atherosclerosis is also present in injured arteries suggests that transitions to plaque VSMC states is an inherent characteristic of VSMCs in different physiological settings. Indeed, there are many commonalities between the two disease models. Both atherosclerotic and injured arteries show endothelial activation at the site of lesions^69,70^. Inflammation is also a key element of both models, with substantial immune cell infiltration into lesions and the underlying media in diseased regions^18,69,71^. Thus, injury and atherosclerosis appear to have an overlap in triggering mechanisms. A notable difference is the absence of high cholesterol in the injury model, contrasting atherosclerosis. However, recent evidence has suggested that plaque cholesterol induces VSMC responses similar to those of inflammation^72^. Perhaps as a consequence, regulation of VSMCs is remarkably comparable in vascular disease models. VSMC proliferation, migration and oligoclonal contribution to lesions is observed in atherosclerosis, aneurysm and injury, and the substantial overlap in gene regulation highlighted here has also been found in other studies^12-16^. For example, a recent scRNA-seq analysis indicated that imVSMCs are present in both atherosclerotic and injured arteries^45^. Our observation that an fcVSMC population is present in both models indicates that also aspects of fibrous cap formation are analogous to VSMC redifferentiation in injury-induced lesions. Importantly, this implies that the carotid ligation injury model, which is acute and has reproducible kinetics compared to experimental atherosclerosis, could be used to model the generation of fibrous cap cells. For instance, manipulation of candidate regulators, such as PAR1, could be used to screen factors for effects on the proportion of fcVSMCs in neointimal lesions at well-characterised timepoints.

The identification of a novel VSMC transition that is associated with fibrous cap formation provides exciting new insight into the VSMC mechanisms underpinning plaque stability. We propose that deeper understanding of this transition and its regulation could provide new drug targets that could be used for promoting fibrous cap formation and thus mitigation of plaque rupture.

## Funding

This work was supported by grants from the British Heart Foundation (PG/19/6/34153, FS/15/62/32032, FS/15/38/31516, RM/13/3/30159, RE/13/6/30180, RE/18/1/34212, CH/2000003/12800 to J.C.K.T., M.D.W., H.F.J., M.R.B.), the NIHR Cambridge Biomedical Research Centre and the Chan Zuckerberg Initiative (2018-190766/RG98793, the Collaborative Bioresource for Translational Medicine, K.T.M.).

## Authors’ contributions

J.C.K.T. and H.F.J. conceived the study. J.C.K.T., M.D.W., J.L., S.O. performed experiments and data analysis. N.L.F., K.F., A.F. and Y.-H.C. performed surgery and histology supervised by M.R.B. and M.C.H. C. K.T.M. retrieved human tissue samples. J.C.K.T. and H.F.J. wrote the manuscript with input from all authors.

## Acknowledgements

We acknowledge the National Institute for Health Research Cambridge Biomedical Research Centre Cell Phenotyping Hub for cell sorting, Darran Clements at the MRC/Wellcome Stem Cell Institute Advanced Imaging Facility for assistance with confocal imaging, the Babraham Institute Sequencing Facility (UKRI-BBSRC Core Capability Grant), Simon Andrews and Felix Krueger (Babraham Institute’s Bioinformatics) for data processing with CellRanger, Lauren Kitt for assistance with animal models. Data analysis was done using the High Performance Computer clusters at the University of Cambridge and Imperial College Research Computing Service (DOI: 10.14469/hpc/2232). We thank the donors, their families, the specialist nurses on organ donation (SNOD), and the Papworth Tissue Bank team for human tissue samples. The Visium experiment was done by the Spatially Resolved Transcriptomics (SpaRTAN) group at Newcastle University.

## Conflict of Interest

H.F.J is a key opinion leader for Novo Nordisk A/S.

## References

1 Bennett, M. R., Sinha, S. & Owens, G. K. Circ Res 118, 692–702, doi:10.1161/circresaha.115.306361 (2016).

2 Basatemur, G. L., Jørgensen, H. F., Clarke, M. C. H., Bennett, M. R. & Mallat, Z. Nat Rev Cardiol 16, 727–744, doi:10.1038/s41569-019-0227-9 (2019).

3 Alexander, M. R. & Owens, G. K. Annu Rev Physiol 74, 13–40, doi:10.1146/annurev-physiol-012110-142315 (2012).

4 Nemenoff, R. A., Horita, H., Ostriker, A. C., Furgeson, S. B., Simpson, P. A., VanPutten, V., Crossno, J., Offermanns, S. & Weiser-Evans, M. C. Arterioscler Thromb Vasc Biol 31, 1300–1308, doi:10.1161/atvbaha.111.223701 (2011).

5 Yang, P., Hong, M. S., Fu, C., Schmit, B. M., Su, Y., Berceli, S. A. & Jiang, Z. Surgery 159, 602–612, doi:10.1016/j.surg.2015.08.015 (2016).

6 Herring, B. P., Hoggatt, A. M., Burlak, C. & Offermanns, S. Vasc Cell 6, 21, doi:10.1186/2045-824x-6-21 (2014).

7 Wang, Z., Wang, D.-Z., Pipes, G. C. T. & Olson, E. N. PNAS 100, 7129, doi:10.1073/pnas.1232341100 (2003).

8 Long, X., Bell, R. D., Gerthoffer, W. T., Zlokovic, B. V. & Miano, J. M. Arterioscler Thromb Vasc Biol 28, 1505–1510, doi:10.1161/atvbaha.108.166066 (2008).

9 Gomez, D. & Owens, G. K. Cardiovasc Res 95, 156–164, doi:10.1093/cvr/cvs115 (2012).

10 Gomez, D., Shankman, L. S., Nguyen, A. T. & Owens, G. K. Nat Methods 10, 171–177, doi:10.1038/nmeth.2332 (2013).

11 Shankman, L. S., Gomez, D., Cherepanova, O. A., Salmon, M., Alencar, G. F., Haskins, R. M., Swiatlowska, P., Newman, A. A., Greene, E. S., Straub, A. C., Isakson, B., Randolph, G. J. & Owens, G. K. Nat Med 21, 628–637, doi:10.1038/nm.3866 (2015).

12 Chappell, J., Harman, J. L., Narasimhan, V. M., Yu, H., Foote, K., Simons, B. D., Bennett, M. R. & Jørgensen, H. F. Circ Res 119, 1313–1323, doi:10.1161/circresaha.116.309799 (2016).

13 Feil, S., Fehrenbacher, B., Lukowski, R., Essmann, F., Schulze-Osthoff, K., Schaller, M. & Feil, R. Circ Res 115, 662–667, doi:10.1161/circresaha.115.304634 (2014).

14 Jacobsen, K., Lund, M. B., Shim, J., Gunnersen, S., Füchtbauer, E.-M., Kjolby, M., Carramolino, L. & Bentzon, J. F. JCI Insight 2, e95890, doi:10.1172/jci.insight.95890 (2017).

15 Misra, A., Feng, Z., Chandran, R. R., Kabir, I., Rotllan, N., Aryal, B., Sheikh, A. Q., Ding, L., Qin, L., Fernández-Hernando, C., Tellides, G. & Greif, D. M. Nat Commun 9, 2073, doi:10.1038/s41467-018-04447-7 (2018).

16 Liu, M. & Gomez, D. Arterioscler Thromb Vasc Biol 39, 1715–1723, doi:10.1161/atvbaha.119.312131 (2019).

17 Dobnikar, L., Taylor, A. L., Chappell, J., Oldach, P., Harman, J. L., Oerton, E., Dzierzak, E., Bennett, M. R., Spivakov, M. & Jørgensen, H. F. Nat Commun 9, 4567, doi:10.1038/s41467-018-06891-x (2018).

18 Worssam, M. D., Lambert, J., Oc, S., Taylor, J. C. K., Taylor, A. L., Dobnikar, L., Chappell, J., Harman, J. L., Figg, N. L., Finigan, A., Foote, K., Uryga, A. K., Bennett, M. R., Spivakov, M. & Jørgensen, H. F. Cardiovasc Res 119, 1279–1294, doi:10.1093/cvr/cvac138 (2023).

19 Conklin, A. C., Nishi, H., Schlamp, F., Örd, T., Õunap, K., Kaikkonen, M. U., Fisher, E. A. & Romanoski, C. E. Immunometabolism 3, e210022, doi:10.20900/immunometab20210022 (2021).

20 Pan, H., Xue, C., Auerbach, B. J., Fan, J., Bashore, A. C., Cui, J., Yang, D. Y., Trignano, S. B., Liu, W., Shi, J., Ihuegbu, C. O., Bush, E. C., Worley, J., Vlahos, L., Laise, P., Solomon, R. A., Connolly, E. S., Califano, A., Sims, P. A., Zhang, H., Li, M. & Reilly, M. P. Circulation 142, 2060–2075, doi:10.1161/CIRCULATIONAHA.120.048378 (2020).

21 Alencar, G. F., Owsiany, K. M., Karnewar, S., Sukhavasi, K., Mocci, G., Nguyen, A. T., Williams, C. M., Shamsuzzaman, S., Mokry, M., Henderson, C. A., Haskins, R., Baylis, R. A., Finn, A. V., McNamara, C. A., Zunder, E. R., Venkata, V., Pasterkamp, G., Björkegren, J., Bekiranov, S. & Owens, G. K. Circulation 142, 2045–2059, doi:10.1161/CIRCULATIONAHA.120.046672 (2020).

22 Newby, A. C. & Zaltsman, A. B. Cardiovasc Res 41, 345–360, doi:10.1016/S0008-6363(98)00286-7 (1999).

23 Rekhter, M. D. Cardiovasc Res 41, 376–384, doi:10.1016/s0008-6363(98)00321-6 (1999).

24 Newman, A. A. C., Serbulea, V., Baylis, R. A., Shankman, L. S., Bradley, X., Alencar, G. F., Owsiany, K., Deaton, R. A., Karnewar, S., Shamsuzzaman, S., Salamon, A., Reddy, M. S., Guo, L., Finn, A., Virmani, R., Cherepanova, O. A. & Owens, G. K. Nat Metab 3, 166–181, doi:10.1038/s42255-020-00338-8 (2021).

25 Wirka, R. C., Wagh, D., Paik, D. T., Pjanic, M., Nguyen, T., Miller, C. L., Kundu, R., Nagao, M., Coller, J., Koyano, T. K., Fong, R., Woo, Y. J., Liu, B., Montgomery, S. B., Wu, J. C., Zhu, K., Chang, R., Alamprese, M., Tallquist, M. D., Kim, J. B. & Quertermous, T. Nat Med 25, 1280–1289, doi:10.1038/s41591-019-0512-5 (2019).

26 Martos-Rodríguez, C. J., Albarrán-Juárez, J., Morales-Cano, D., Caballero, A., MacGrogan, D., de la Pompa, J. L., Carramolino, L. & Bentzon, J. F. Arteriosclerosis, Thrombosis, and Vascular Biology 0, ATVBAHA.120.315627, doi:10.1161/ATVBAHA.120.315627 (2021).

27 Hafemeister, C. & Satija, R. Genom Biol 20, 296, doi:10.1186/s13059-019-1874-1 (2019).

28 Satija, R., Farrell, J. A., Gennert, D., Schier, A. F. & Regev, A. Nature Biotechnology 33, 495–502, doi:10.1038/nbt.3192 (2015).

29 Ovchinnikova, S. & Anders, S. Genome Res 30, 749–756, doi:10.1101/gr.251447.119 (2020).

30 Street, K., Risso, D., Fletcher, R. B., Das, D., Ngai, J., Yosef, N., Purdom, E. & Dudoit, S. BMC Genomics 19, 477, doi:10.1186/s12864-018-4772-0 (2018).

31 Lun, A. & D., R. (Bioconductor, https://bioconductor.org/packages/release/bioc/html/SingleCellExperiment.html, 2020).

32 Hastie, T. (CRAN, https://cran.r-project.org/web/packages/gam/index.html, 2019).

33 Kolde, R. (CRAN, https://cran.r-project.org/web/packages/pheatmap/index.html, 2019).

34 Wolf, F. A., Hamey, F. K., Plass, M., Solana, J., Dahlin, J. S., Göttgens, B., Rajewsky, N., Simon, L. & Theis, F. J. Genom Biol 20, 59, doi:10.1186/s13059-019-1663-x (2019).

35 Wolf, F. A., Angerer, P. & Theis, F. J. Genom Biol 19, 15, doi:10.1186/s13059-017-1382-0 (2018).

36 Philippe, A. (Pypi, https://pypi.org/project/anndata2ri/#description, 2020).

37 Raudvere, U., Kolberg, L., Kuzmin, I., Arak, T., Adler, P., Peterson, H. & Vilo, J. Nucleic Acid Res 47, W191–W198, doi:10.1093/nar/gkz369 (2019).

38 Queen, R., Crosier, M., Eley, L., Kerwin, J., Turner, J. E., Yu, J., Alqahtani, A., Dhanaseelan, T., Overman, L., Soetjoadi, H., Baldock, R., Coxhead, J., Boczonadi, V., Laude, A., Cockell, S. J., Kane, M. A., Lisgo, S. & Henderson, D. J. PLoS Genet 19, e1010777, doi:10.1371/journal.pgen.1010777 (2023).

39 Ritchie, M. E., Phipson, B., Wu, D., Hu, Y., Law, C. W., Shi, W. & Smyth, G. K. Nucleic Acids Res 43, e47, doi:10.1093/nar/gkv007 (2015).

40 Alonso-Herranz, L., Albarrán-Juárez, J. & Bentzon, J. F. Front Cardiovasc Med 10, 1254114, doi:10.3389/fcvm.2023.1254114 (2023).

41 Lu, Y. W., Lowery, A. M., Sun, L. Y., Singer, H. A., Dai, G., Adam, A. P., Vincent, P. A. & Schwarz, J. J. Arterioscler Thromb Vasc Biol 37, 1380–1390, doi:10.1161/atvbaha.117.309180 (2017).

42 Sharma, D., Worssam, M. D., Pedroza, A. J., Dalal, A. R., Alemany, H., Kim, H. J., Kundu, R., Fischbein, M. P., Cheng, P., Wirka, R. & Quertermous, T. Arterioscler Thromb Vasc Biol 44, 391–408, doi:10.1161/atvbaha.123.320030 (2024).

43 Kumar, A. & Lindner, V. Arterioscler Thromb Vasc Biol 17, 2238–2244, doi:10.1161/01.ATV.17.10.2238 (1997).

44 Regan, C. P., Adam, P. J., Madsen, C. S. & Owens, G. K. J Clin Invest 106, 1139–1147, doi:10.1172/jci10522 (2000).

45 Warwick, T., Buchmann, G. K., Pflüger-Müller, B., Spaeth, M., Schürmann, C., Abplanalp, W., Tombor, L., John, D., Weigert, A., Leo-Hansmann, M., Dimmeler, S. & Brandes, R. P. Front Physiol 14, 1125864, doi:10.3389/fphys.2023.1125864 (2023).

46 Jansen, I., Cahalane, R., Hengst, R., Akyildiz, A., Farrell, E., Gijsen, F., Aikawa, E., van der Heiden, K. & Wissing, T. Basic Res Cardiol 119, 193–213, doi:10.1007/s00395-024-01033-5 (2024).

47 Schrör, K., Bretschneider, E., Fischer, K., Fischer, J. W., Pape, R., Rauch, B. H., Rosenkranz, A. C. & Weber, A. A. Thromb Haemost 103, 884–890, doi:10.1160/th09-09-0627 (2010).

48 Fager, G. Circ Res 77, 645–650, doi:doi:10.1161/01.RES.77.4.645 (1995).

49 Cicha, I., Yilmaz, A., Klein, M., Raithel, D., Brigstock, D. R., Daniel, W. G., Goppelt-Struebe, M. & Garlichs, C. D. Arterioscler Thromb Vasc Biol 25, 1008–1013, doi:10.1161/01.ATV.0000162173.27682.7b (2005).

50 Wang, Y., Liu, X., Xu, Q., Xu, W., Zhou, X., Leask, A. & Lin, Z. JCI Insight 8, doi:10.1172/jci.insight.162987 (2023).

51 Karnik, S. K., Brooke, B. S., Bayes-Genis, A., Sorensen, L., Wythe, J. D., Schwartz, R. S., Keating, M. T. & Li, D. Y. Development 130, 411–423, doi:10.1242/dev.00223 (2003).

52 Moura, R., Tjwa, M., Vandervoort, P., Van Kerckhoven, S., Holvoet, P. & Hoylaerts, M. F. Circ Res 103, 1181–1189, doi:10.1161/circresaha.108.185645 (2008).

53 de Waard, V., van Achterberg, T. A., Beauchamp, N. J., Pannekoek, H. & de Vries, C. J. Arterioscler Thromb Vasc Biol 23, 64–68, doi:10.1161/01.atv.0000042218.13101.50 (2003).

54 Fouillade, C., Monet-Leprêtre, M., Baron-Menguy, C. & Joutel, A. Cardiovasc Res 95, 138–146, doi:10.1093/cvr/cvs019 (2012).

55 Tang, Y., Urs, S., Boucher, J., Bernaiche, T., Venkatesh, D., Spicer, D. B., Vary, C. P. & Liaw, L. J Biol Chem 285, 17556–17563, doi:10.1074/jbc.M109.076414 (2010).

56 Janjanam, J., Zhang, B., Mani, A. M., Singh, N. K., Traylor, J. G., Jr., Orr, A. W. & Rao, G. N. J Biol Chem 293, 3088–3103, doi:10.1074/jbc.RA117.000866 (2018).

57 Ko, W. C., Chen, B. C., Hsu, M. J., Tsai, C. T., Hong, C. Y. & Lin, C. H. Acta Pharmacol Sin 33, 49–56, doi:10.1038/aps.2011.178 (2012).

58 Jaberi, N., Soleimani, A., Pashirzad, M., Abdeahad, H., Mohammadi, F., Khoshakhlagh, M., Khazaei, M., Ferns, G. A., Avan, A. & Hassanian, S. M. J Cell Biochem 120, 4757–4765, doi:10.1002/jcb.27771 (2019).

59 Kalz, J., ten Cate, H. & Spronk, H. M. J Thromb Thrombolysis 37, 45–55, doi:10.1007/s11239-013-1026-5 (2014).

60 Pingel, S., Tiyerili, V., Mueller, J., Werner, N., Nickenig, G. & Mueller, C. Arch Med Sci 10, 154–160, doi:10.5114/aoms.2014.40742 (2014).

61 Spronk, H. M. H., Borissoff, J. I., Soehnlein, O., Koenen, R., van Oerle, R., Heeneman, S., Daemen, M. J. A. P., Esmon, C. T., Degen, J. L., Weiler, H., van Ryn, J. & Cate, H. T. Blood 120, 103, doi:10.1182/blood.V120.21.103.103 (2012).

62 Friebel, J., Moritz, E., Witkowski, M., Jakobs, K., Strässler, E., Dörner, A., Steffens, D., Puccini, M., Lammel, S., Glauben, R., Nowak, F., Kränkel, N., Haghikia, A., Moos, V., Schutheiss, H. P., Felix, S. B., Landmesser, U., Rauch, B. H. & Rauch, U. Cells 10, doi:10.3390/cells10123517 (2021).

63 Dollery, C. M. & Libby, P. Cardiovascular Research 69, 625–635, doi:10.1016/j.cardiores.2005.11.003 (2006).

64 Heuberger, D. M. & Schuepbach, R. A. Thrombosis Journal 17, 4, doi:10.1186/s12959-019-0194-8 (2019).

65 Lee, C. J. & Ansell, J. E. Br J Clin Pharmacol 72, 581–592, doi:10.1111/j.1365-2125.2011.03916.x (2011).

66 Tantry, U. S., Bliden, K. P., Chaudhary, R., Novakovic, M., Rout, A. & Gurbel, P. A. Future Cardiol 16, 373–384, doi:10.2217/fca-2019-0090 (2020).

67 van Dam-Nolen, D. H. K., Truijman, M. T. B., van der Kolk, A. G., Liem, M. I., Schreuder, F., Boersma, E., Daemen, M., Mess, W. H., van Oostenbrugge, R. J., van der Steen, A. F. W., Bos, D., Koudstaal, P. J., Nederkoorn, P. J., Hendrikse, J., van der Lugt, A. & Kooi, M. E. JACC Cardiovasc Imaging 15, 1715–1726, doi:10.1016/j.jcmg.2022.04.003 (2022).

68 Tian, J., Wang, X., Tian, J. & Yu, B. BMC Cardiovasc Disord 19, 45, doi:10.1186/s12872-019-1023-5 (2019).

69 Ding, X., An, Q., Zhao, W., Song, Y., Tang, X., Wang, J., Chang, C. C., Zhao, G., Hsiai, T., Fan, G., Fan, Y. & Li, S. JCI Insight 7, doi:10.1172/jci.insight.153769 (2022).

70 Gimbrone, M. A., Jr. & García-Cardeña, G. Circ Res 118, 620–636, doi:10.1161/circresaha.115.306301 (2016).

71 Wolf, D. & Ley, K. Circ Res 124, 315–327, doi:10.1161/circresaha.118.313591 (2019).

72 Carramolino, L., Albarrán-Juárez, J., Markov, A., Hernández-SanMiguel, E., Sharysh, D., Cumbicus, V., Morales-Cano, D., Labrador-Cantarero, V., Møller, P. L., Nogales, P., Benguria, A., Dopazo, A., Sanchez-Cabo, F., Torroja, C. & Bentzon, J. F. Nature Cardiovascular Research 3, 203–220, doi:10.1038/s44161-023-00412-w (2024).

