## Supplemental Information for "Delineation of a thrombin receptor-stimulated vascular smooth muscle cell transition generating cells in the plaque-stabilising fibrous cap"

Short title: Thrombin receptor induces a fibrous cap VSMC state

Content:

Supplementary Table 1: scRNA-seq data analysis parameters (**see separate spreadsheet)**

Supplementary Table 2: RT-qPCR primer sequences

Supplementary Table 3: GO-term analysis of fcVSMC vs imVSMC upregulated genes in mouse atherosclerosis (**see separate spreadsheet)**

Supplementary Table 3: GO-term analysis of thrombin-treated vs control hVSMC upregulated genes (**see separate spreadsheet)**

Supplementary Figure 1: Atherosclerosis dataset split by vessel health status and HFD timepoint

Supplementary Figure 2: Injury scRNA-seq dataset plot showing Notch3 expression

Supplementary Figure 3: Partition-based graph abstraction (PAGA) analysis of the injury dataset

Supplementary Figure 4: Quantification of medial VCAM1 and NOTCH3 staining following injury

Supplementary Figure 5: Dot plot of Visium dataset showing cell state markers across clusters

Supplementary Figure 6: Gene expression following thrombin treatment of hVSMCs with and without the PAR1 inhibitor, vorapaxar.

### Supplementary Tables

Supplementary Tables 1, 3, 4 are provided as separate files

**Supplementary Table 2: RT-qPCR primer sequences**

| **Gene** | **Forward Primer** | **Reverse Primer** |
|---|---|---|
| HMBS | GGTTGTTCACTCCTTGAAGG | TTCTCTGGCAGGGTTTCTAG |
| ACTA2 | AGATCTGGCACCACTCTTTC | GTGAGTCACACCATCTCCAG |
| CNN1 | CCAACGACCTGTTTGAGAACACC | ATTTCCGCTCCTGCTTCTCTGC |
| TAGLN | TCTTTGAAGGCAAAGACATGG | TTATGCTCCTGCGCTTTCTT |
| CCN2 | CTTGCGAAGCTGACCTGGAAGA | CCGTCGGTACATACTCCACAGA |
| ELN | GGTTGTGTCACCAGAAGCAGCT | CCGTAAGTAGGAATGCCTCCAAC |
| THBS1 | GCTGGAAATGTGGTGCTTGTCC | CTCCATTGTGGTTGAAGCAGGC |
| ANKRD1 | CGACTCCTGATTATGTATGGCGC | GCTTTGGTTCCATTCTGCCAGTG |

### Supplementary Figures

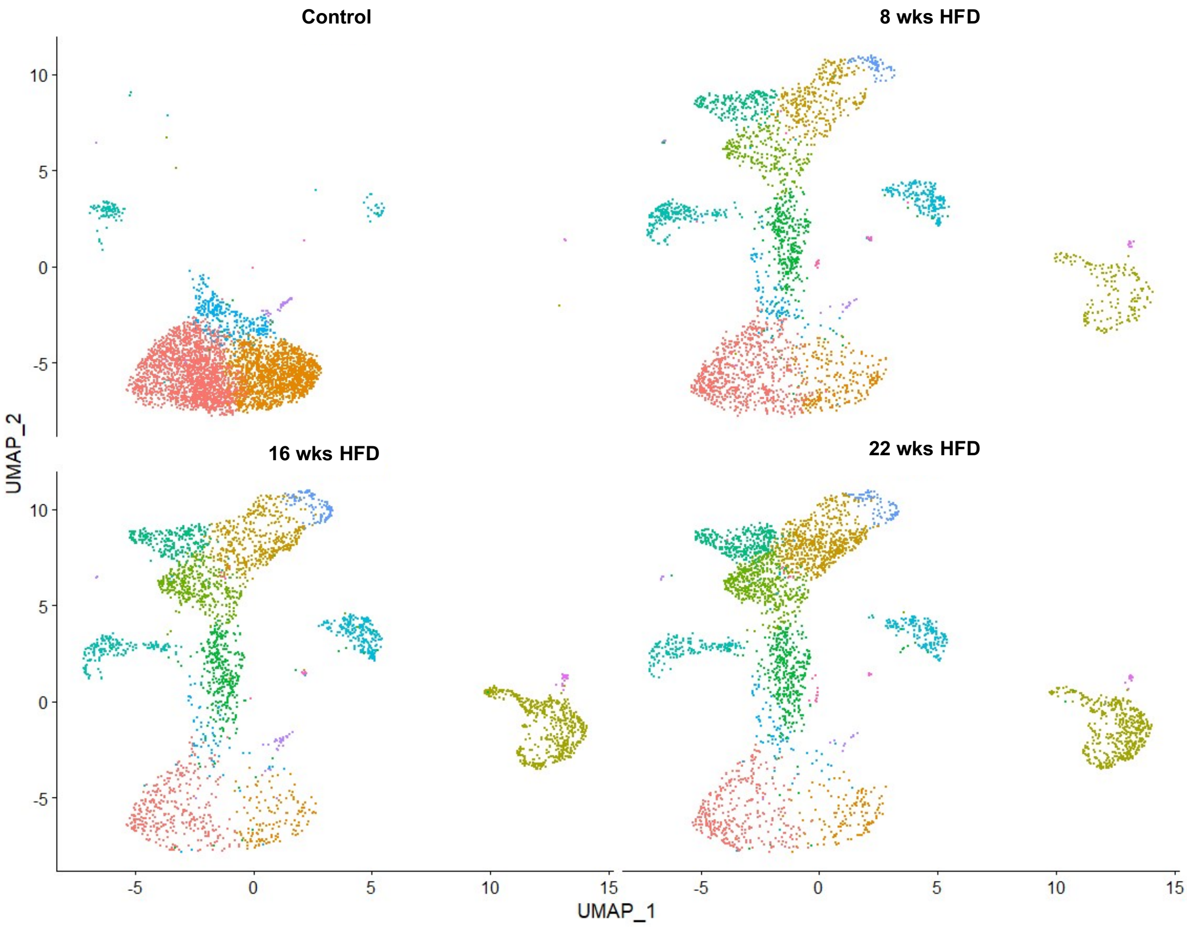


**Supplementary Figure 1. Atherosclerosis scRNA-seq dataset split by vessel health status and HFD timepoint.** Uniform Manifold Approximation and Projection (UMAP) with cell cluster map separated by vessel health status/high fat diet (HFD) timepoint for scRNA-seq analysis of VSMC-lineage label positive cells from atherosclerotic arteries (GSE155513) taken from Myh11-ZsGreen/ApoE-/- animals after 8, 16 and 22 weeks of HFD feeding and healthy control Apoe$^{+/+}$ carotid arteries.

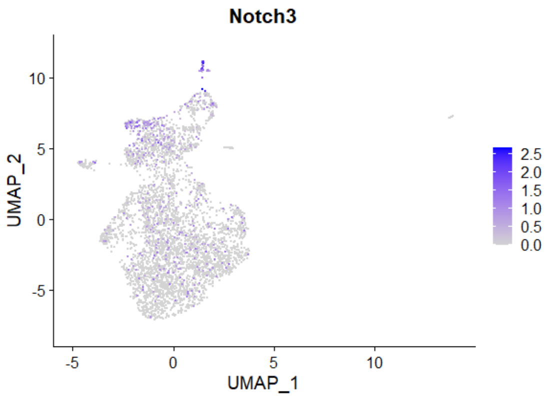


**Supplementary Figure 2. Notch3 expression in VSMCs 11 days after injury.** Uniform Manifold Approximation and Projection (UMAP) of VSMC-lineage label positive cells isolated from left carotid arteries 11 days after carotid ligation injury (11 DPI) showing *Notch3* expression with a grey (low) to blue (high) scale.

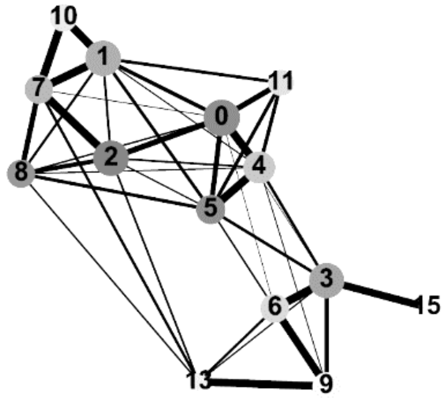


**Supplementary Figure 3. Partition-based graph abstraction (PAGA) analysis of the injury dataset.** PAGA plot showing cluster connectivity in the injury dataset of VSMC-lineage label positive cells sorted from left carotid arteries 11 days after carotid ligation injury (11 DPI). Line thickness indicates degree of transcriptional similarity between clusters.

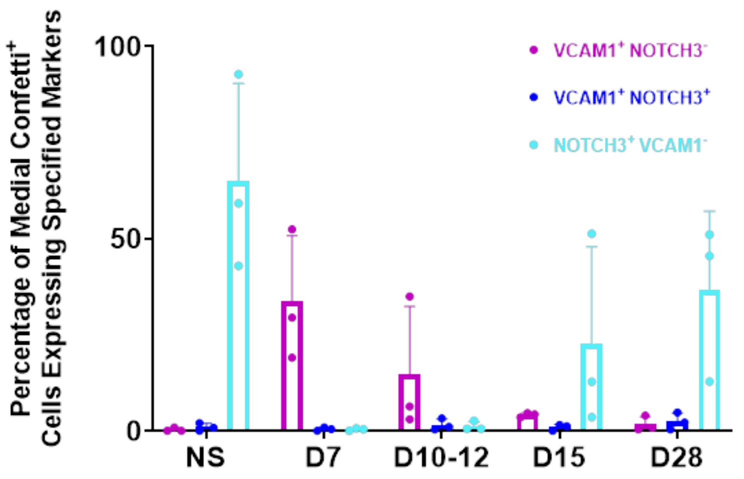


**Supplementary Figure 4. Quantification of medial VCAM1 and NOTCH3 staining following injury.** The proportion of medial Confetti+ cells in diseased artery regions with specified marker expression at indicated timepoints after injury (n=3 animals per timepoint).

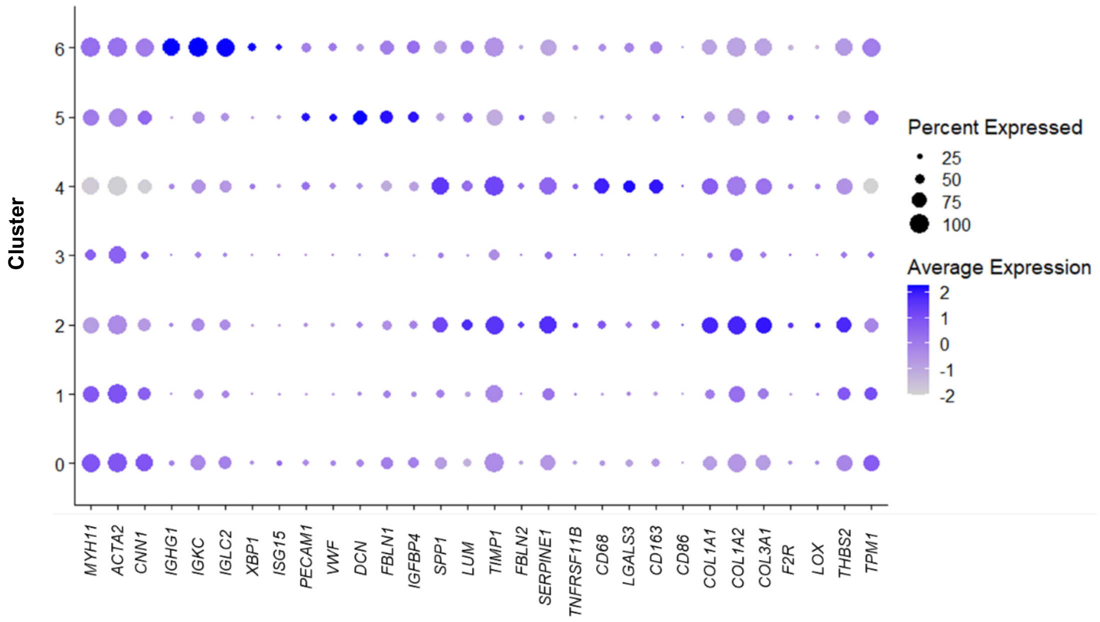


**Supplementary Figure 5. Human plaque spatial transcriptomic dataset dot plot showing cell state markers across clusters.** Dot plot showing expression of cell type markers in clusters using a scale from grey (low) to purple (high). The percentage of capture regions within clusters with detected marker expression is indicated by dot size.

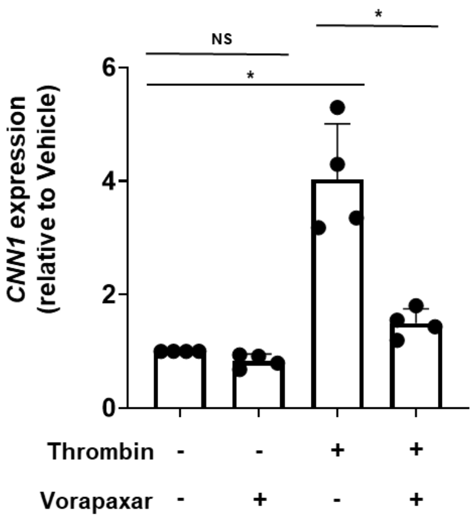


**Supplementary Figure 6. Gene expression effect following thrombin treatment of hVSMCs with and without the PAR1 inhibitor, vorapaxar.** Expression levels of CNN1 in thrombin-treated human VSMCs relative to vehicle-treated samples with and without co-treatment with vorapaxar. Bars show mean ± SD. n=4 cell lines (female). *P-adj<0.05, NS: not significant (Mann-Whitney with Bonferroni multiple testing correction, Statistical testing was done between groups).
